# Spatially-resolved transcriptomics analyses of invasive fronts in solid tumors

**DOI:** 10.1101/2021.10.21.465135

**Authors:** Liang Wu, Jiayan Yan, Yinqi Bai, Feiyu Chen, JiangShan Xu, Xuanxuan Zou, Ao Huang, Liangzhen Hou, Yu Zhong, Zehua Jing, Xiaorui Zhou, Haixiang Sun, Mengnan Cheng, Yuan Ji, Rongkui Luo, Qinqin Li, Liang Wu, Pengxiang Wang, Dezhen Guo, Waidong Huang, Junjie Lei, Sha Liao, Yuxiang Li, Zhifeng Jiang, Na Yao, Yang Yu, Yao Li, Fengming Liu, Mingyuan Zhang, Huanming Yang, Shuang Yang, Xun Xu, Longqi Liu, Xiangdong Wang, Jian Wang, Jia Fan, Shiping Liu, Xinrong Yang, Ao Chen, Jian Zhou

## Abstract

Solid tumors are complex ecosystems, and heterogeneity is the major challenge for overcoming tumor relapse and metastasis. Uncovering the spatial heterogeneity of cell types and functional states in tumors is essential for developing effective treatment, especially in invasive fronts of tumor, the most active region for tumor cells infiltration and invasion. We firstly used SpaTial Enhanced REsolution Omics-sequencing (Stereo-seq) with a nanoscale resolution to characterize the tumor microenvironment of intrahepatic cholangiocarcinoma (ICC). Enrichment of distinctive immune cells, suppressive immune microenvironment and metabolic reprogramming of tumor cells were identified in the 500µm-wide zone centered bilaterally on the tumor boundary, namely invasive fronts of tumor. Furthermore, we found the damaged states of hepatocytes with overexpression of Serum Amyloid A (SAA) in invasive fronts, recruiting macrophages for facilitating further tumor invasion, and thus resulting in a worse prognosis. We also confirmed these findings in hepatocellular carcinoma and other liver metastatic cancers. Our work highlights the remarkable potential of high-resolution-spatially resolved transcriptomic approaches to provide meaningful biological insights for comprehensively dissecting the tumor ecosystem and guiding the development of novel therapeutic strategies for solid tumors.

---

Solid tumors are complex organ-like structures, which tend to spread to nearby tissues and metastasize to remote organs through the lymph system and bloodstream^1–4^. Evidences have indicated that solid tumors are complex ecosystems with a high degree of heterogeneity, involving areas where cancer cells interact with various cell types like immune cells and stromal cells, as well as extracellular matrix (ECM) components^5, 6^. Specifically, tumor invasive margin area, where tumor cells invade into paranormal tissues and encountered a diverse array of stromal cell and ECM components, which was the most active region for tumor cells infiltration and invasion^4, 7–11^. A complete understanding of the components and their spatial heterogeneities, and the interplay between tumor cells and tumor microenvironment (TME) in this area, will facilitate to understand tumor invasion and metastasis, evaluate the prognoses, and explore novel therapeutic approaches for solid tumors^5, 6, 11–14^.

Single-cell RNA sequencing (scRNA-seq) is a powerful tool for the investigation of cellular components and their interactions at the single-cell level, which has been used to characterize the TME of several types of solid tumors^15, 16^. However, the lack of spatial information is a major obstacle in interrogating the correlations between the local environment, specific cell-cell interactions, and tumor progression, especially in tumor invasive margin area. Furthermore, due to the lack of multi-regional sampling in previous studies, intratumoral spatial heterogeneity at the resolution of single cells remains poorly characterized^17–19^. The recently developed spatial transcriptomics (ST) method facilitates unbiased mapping of transcripts over entire tissue sections using spatially barcoded oligo-deoxythymidine microarrays^20, 21^. However, the majority of ST approaches applied in previous studies were of a low resolution, recognizing spots mixed with dozens of cells instead of single cells^12, 13^. Recently, we combined DNA nanoball (DNB) patterned array chips and *in situ* RNA capture to develop SpaTial Enhanced REsolution Omics-sequencing (Stereo-seq) with the nanoscale resolution (220 nm × 220 nm/spot) and expandable areas (10 mm × 10 mm) with a few hundred spots of data captured per cell^12^. Stereo-seq can therefore potentially bridge the gap between scRNA-seq and ST analyses, which together can better characterize functional and structural studies of entire tumor ecosystems. Thus, integration of single-cell and high dimensional spatial data produced by Stereo-seq from tumor multi-regional tissues would facilitate comprehensive and unbiased tissue analyses to identify intratumor heterogeneities and TME cellular communications in solid tumor, especially for tumor invasive margin area.

Liver cancer is one of the most malignant solid tumors worldwide, while intrahepatic cholangiocarcinoma (ICC) is the second most common primary liver cancer with an increasing global incidence during past decades^22–24^. However, most patients (> 70%) are already at advanced stages at the time of diagnoses and cannot be surgically treated due to locally advanced or metastatic disease^25–27^. In this study, we used Stereo-seq with high resolution and centimeter-sized fields of view to characterize the complexity and heterogeneity of tumor ecosystems, as well as their cellular interactions in ICC, by analyzing four regional sites, including tumor tissue (T), margin areas (B), adjacent normal tissue (P), and normal or metastatic lymph nodes (LN). By integrating ST data with scRNA-seq data and using bioinformatics analyses, our approach determined a high degree of cellular and transcriptional heterogeneities in the tumor invasive front, where tumor cells invade into paranormal tissues. We found an increased immune-escape capacity and a shift toward increased fatty metabolism of tumor cells, as well as an enhanced accumulation of immune cells, especially in the region within 250 µm from the tumor-normal borderline. Furthermore, we found increased expressions of *SAA1* and *SAA2* in hepatocytes close to invasive fronts, which were associated with enhanced local recruitment of macrophages and correlated with worse prognoses in ICC patients. Furthermore, we confirmed these findings in primary and metastatic liver cancer with other four additional cohorts. Our study highlighted the important potential of high resolution, spatially-resolved transcriptomics approaches in providing meaningful biological insights for the development of novel therapeutic strategies for solid tumors.

## Results

### Spatial resolved transcriptomics profiles of multi-regional tissues from ICC patients

To characterize the spatial transcriptional landscape of ICC, we processed fresh frozen tissues from the intratumor tissues, adjacent normal tissues, margin areas, and normal or metastatic lymph nodes using Stereo-seq with high resolution (220 nm × 220 nm/spot) and expandable areas (10 mm × 10 mm) (Discovery Cohort, Fig. 1a, see Methods). We generated Stereo-seq data of 50 slides from 27 samples (T, 6; B, 8; P, 7; LN, 6) of eight patients pathologically diagnosed with ICC (LC0, LC1, LC2, LC4, LC5, LC6, LC7, and LC8) using hemoxylin and eosin (H&E) staining of adjacent slides (Fig. 1a and supplementary Table 1). Detailed clinical and pathological information is provided in Supplementary Table 2. Moreover, we recruited additional 21 patients with liver cancer with margin areas stained by multiplexed immunofluorescence (IF) (Validation Cohort 1), 10 ICC patients with matched tumor tissues, border areas and adjacent normal tissues for bulk-RNA sequencing (Validation Cohort 2), and 93 ICC patients with paired tumor tissues and adjacent normal tissues for immunohistochemistry (IHC) staining (Validation Cohort 3). Additionally, 159 HCC patients with paired tumor tissues and paratumor tissues for bulk-RNA sequencing and proteomics recruited in our previous study^28^ were used as Validation Cohort 4 (Fig. 1a, Supplementary Table 3). In the Discovery Cohort, scRNA-seq data generated from margin area tissues of patient LC5 (604 cells) were integrated with a recently published ICC scRNA-seq dataset^29^ to establish a reference expression fingerprint of cell types, to delineate the spatial tomographies of cell type populations in each tissue slide (Fig. 1a). As a result, 29,793 qualified cells were clustered into 11 main cell types using Seurat^30^, including T cells (*CD3D*), natural killer cells (NKs) (*KLRF1*), B cells (*MS4A1*), plasma cells (*MZB1*), macrophages (*CD163/CD14*), dendritic cells (DCs) (*CD1C*), cholangiocytes/cholangiocarcinoma cells (*KRT19/EPCAM*), hepatocytes (*ALB*), endothelial cells (*CDH5/ENG*), and fibroblasts (*ACTA2*) (Extended Data Fig. 1a, b). For ST, 2∼90 (a median of eight) transcripts were detected for each DNB or bin (220 nm × 220 nm) and there were about 2,500 (50 × 50) bins for each hepatocyte (with diameters of 25∼30 µm) and about 900 (30 × 30) bins for each cholangiocarcinoma cell (with a diameter of 15 µm) (Extended Data Fig. 1c). To enable spatial cell type annotations with these reference signatures, the raw spatial expression matrix was converted into pseudo-spots with 25 µm square sizes (50 × 50 bins/spot), which approximately represented one cell, and an average of 718 genes and 1,644 mRNA molecules were detected per spot (Extended Data Fig. 1d, e). Cell components for each spot (50 × 50 bins/spot, 25 µm squares) in the ST slides were determined by SPOTlight^31^ using scRNA-seq as the reference, then 11 main cell types were annotated in spatial spots (Fig. 1b, see Methods). We assigned each spot to a specific cell type that showed the highest probabilistic proportion as the first probability was much higher than the second probability (Fig. 1c). The distinct higher expressions of classical cell type marker genes in defined cell clusters supported the rationality of cell type annotations for ST spots (Fig. 1d).

**Fig. 1.**
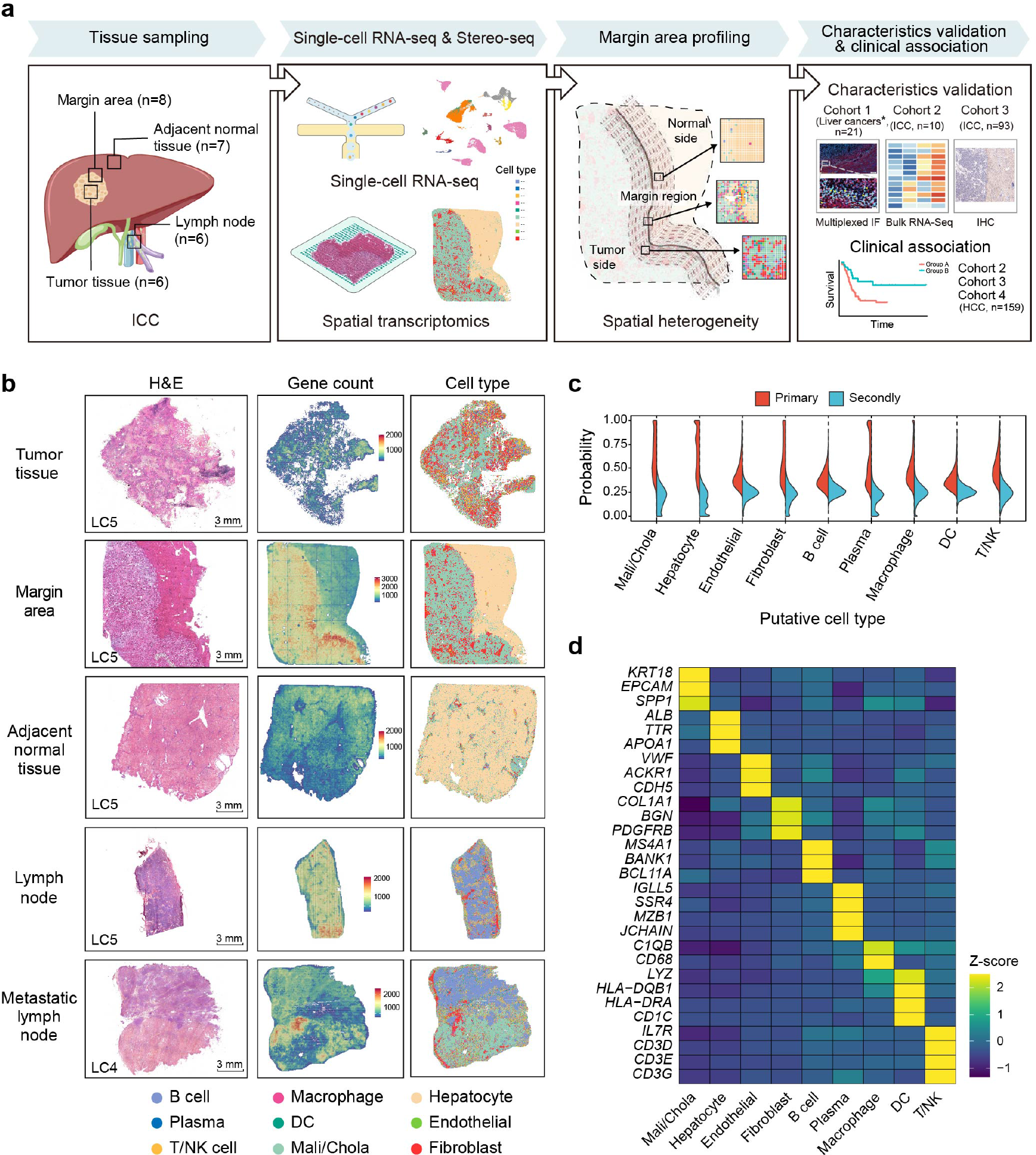
Spatially-resolved transcriptomics profiles in multi-regional sites in patients with intrahepatic cholangiocarcinoma (ICC). (a) Schematic diagram of the spatial transcriptomics acquisition workflow based on 27 samples (T, 6; B, 8; P, 7; LN, 6) from eight patients with ICC with further data analysis. Liver cancers* include ICC (*n* = 10), HCC (*n* = 5), liver metastasis of colorectal cancer (*n* = 3), pancreatic cancer (*n* = 2), and lung cancer (*n* = 1). (b) Hemoxylin and eosin (H&E) staining, gene count maps, and cell type maps of different sites of the ICC patient LC5. (c) The probabilistic inference of cell types at captured spatial transcriptomic spots (50 × 50 bins/spot, 25 µm squares). (d) Heat map showing the expression of marker genes for cell types in annotated spatial spots. T, tumor tissue; B, margin area; P, adjacent normal tissue; LN, lymph node.

### The heterogeneities of cell and ECM compositions as well as spatial distributions in multi-regions of ICC

Cell compositions and spatial distributions were highly heterogeneous in four regional sites (T, P, B, and LN) of patients with ICC. Different areas were spatially characterized with distinct tissue architectures, comprised of different prevailing cell components. In tumor tissues, fibroblasts, macrophages, and B cells were the most abundant cells, which accounted for almost half of all total cell components (Fig. 2a, b). Specifically, a larger percentage of fibroblasts was detected in tumor tissues than in the other three regions, implying the highly desmoplastic property of ICC tumor tissues. As expected, hepatocytes were the prevailing cell type in adjacent normal tissues, followed by cholangiocytes and B cells (Fig. 2a, b). For margin areas, fibroblasts, B cells, and macrophages were the most abundant cell types, except for tumor cells and hepatocytes (Fig. 2a, b). In LNs, B cells, T/NK cells, and macrophages were the most abundant immune cell types, and B cells were scattered widely among the entire LN tissues. Different from normal LNs, in LC4- LN (with LN metastases), tumor cells covered almost half of the LN area in the slide, while B cells covered the remaining normal areas (Fig. 2a, b). Furthermore, typical tertiary lymphoid structures (TLS)^32, 33^, resulting from the co-aggregation of DCs, T/NK, and B cells in non-lymphoid tissues, were also detected in T, P, and B, which was further confirmed by H&E staining (Fig. 2c and Extended Data Fig. 2a). Furthermore, several cell types exhibited distinct enrichments in different tissue areas with high spatial heterogeneities. Most of the fibroblasts in tumor tissues showed matrix cancer-associated fibroblast (mCAF)^29^ signatures with increased expressions of ECM-related genes (*LUM*, *DCN*, and *COL3A1*), while more antigen-presenting cancer-associated fibroblasts (AP-CAF) and inflammatory cancer-associated fibroblasts (iCAF) were found in adjacent normal tissues (Fig. 2d and Extended Data Fig. 2b, c). Compared with adjacent normal tissues, the ratios between exhausted and cytotoxic T cells were higher in tumor tissues and in margin areas (Fig. 2d), indicating suppressive states of T cells and subsequent cancer immune evasion in the TME. After subdividing T cells and B cells into different subpopulations, a significantly larger percentage of naïve T and naïve B cells were observed in LNs, and spatially naïve B cells were mainly surrounded by plasma B cells (Fig. 2d and Extended Data Fig. 2d). Germinal centers characterized by aggregation of DCs surrounded by B cells were also identified in spatial cell type maps (Extended Data Fig. 2d). Apart from the spatial heterogeneous distribution of fibroblast clusters and immune cells in different sites, tumor cells also exhibited distinct enrichment of cancer hallmarks^34^ in different intratumoral areas, indicating spatial heterogeneity between different tissues as well as within tumor tissues (Fig. 2b and Extended Data Fig. 2e).

**Fig. 2.**
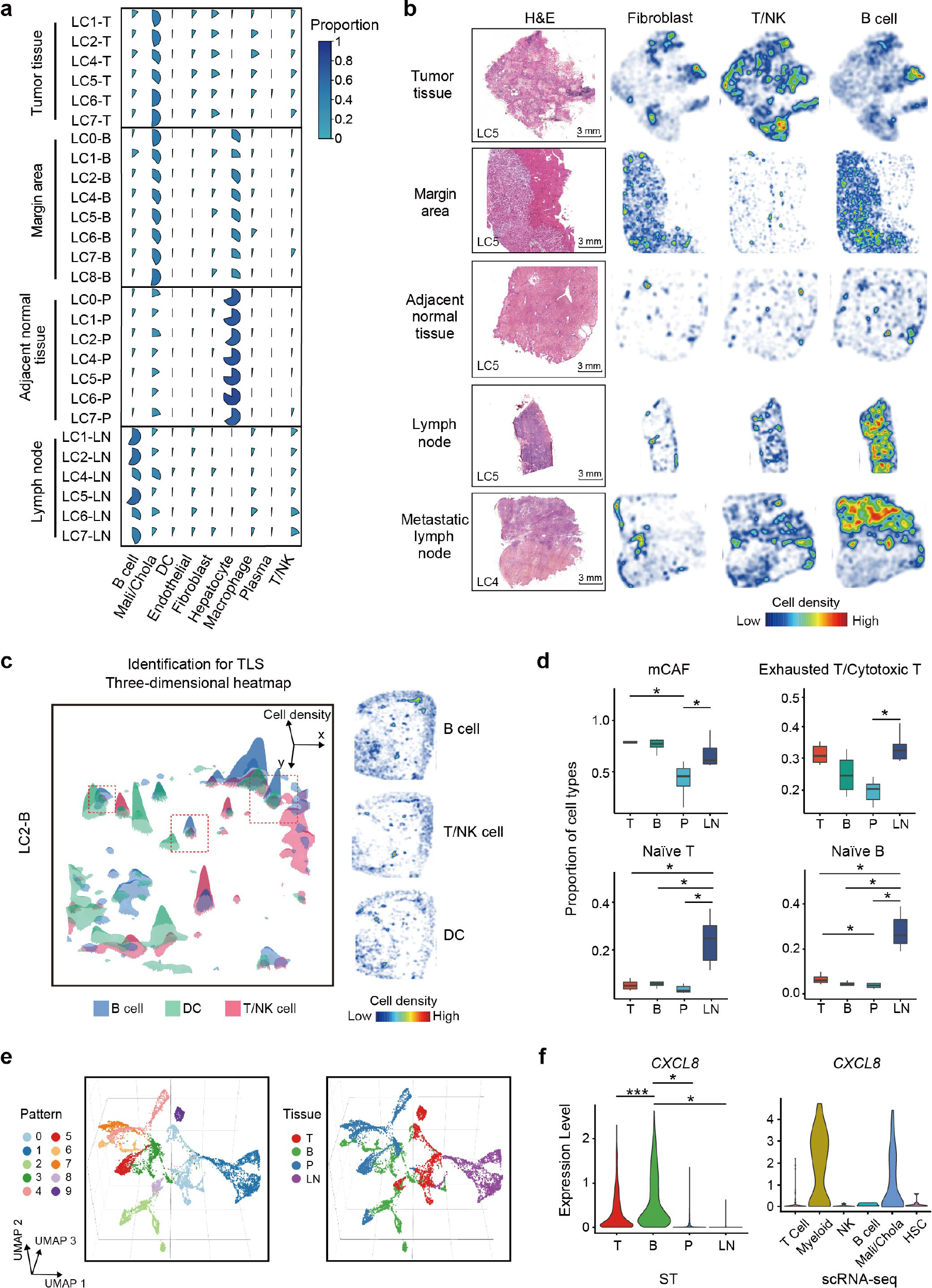
The heterogeneities in composition and spatial distribution of cells and extracellular matrix components in multi-regional sites of ICC patients. **(a)** Pie graphs showing the percentages of major cell types in all cell components at four regional sites (T, *n* = 6; P, *n* = 7; B, *n* = 8; LN, *n* = 6) of eight patients with ICC. **(b)** H&E staining and heat maps of the spatial distributions of fibroblasts, T/NK cells, and B cells in four regional sites. **(c)** The three-dimensional diagram and corresponding heat maps showing tertiary lymphoid structures (TLS) where DCs, T/NK, and B cells co-aggregated. **(d)** Box plots showing the percentages of mCAF, naïve T cells, and naïve B cells in all cell components, and the exhausted/cytotoxic T cells ratios in four regional sites. **(e)** Uniform manifold approximation and projection (UMAP) plots showing the clustering of gene expression profiles of extracellular matrix components in segmented spots (500 µm × 500 µm/spot) from spatial transcriptomics slides. The colors represent different clusters (left panel) and tissues (right panel). **(f)** Violin plots representing the expressions of *CXCL8* in four regional sites for ST (left panel) and main cell types for scRNA-seq (right panel).

The TME is dynamically shaped by bidirectional communication between tumor cells and the ECM through cell-matrix interactions and ECM remodeling. To comprehensively understand ECM remodeling and cell-matrix interactions among four regional sites, we initiated a regional segmentation (1000 × 1000 bins or 500 µm × 500 µm per spot) to obtain approximately 400 spots of local bulk RNA profiles on each ST slide. A total of 20,400 ST spots captured from 48 slides were separated into 10 patterns according to transcriptional signatures of ECM related and chemokines/cytokine genes (Fig. 2e and Extended Data Fig. 3a). Segmented ST spots from the same site gathered from different patients were visualized by uniform manifold approximation and projection (UMAP), indicating a site-specific ECM and chemokine/cytokine enriched microenvironment (Fig. 2e and Extended Data Fig. 3a). In tumor tissues, several highly expressed genes including *HIF1A*, *CXCL6*, *IL18*, and *MMP14* were identified (Extended Data Fig.3a, b). MMP14, which may promote tumor growth and invasion by degrading the matrix barrier and enhancing angiogenesis, was identified as the most enriched MMP in tumors, and was mainly secreted by fibroblasts and tumor cells, based on scRNA-seq data (Extended Data Fig. 3b). Furthermore, *IL18*, encoding a pro-inflammatory cytokine promoting tumor progression^35^, was most abundant in tumors and mainly produced by DCs, macrophages, and tumor cells (Extended Data Fig. 3b). In adjacent normal tissues, ECM components including *IL6R*, *IL1RAP*, and *CXCL2* were highly expressed, when compared with those among the other three areas. The expressions of clusters of chemokines or cytokines, including *CXCL8* and *CXCL1*, were found to be mostly enriched in margin areas, implying strong inflammatory responses in these areas (Fig. 2f and Extended Data Fig. 3b). Specifically, *CXCL8*, encoding a major mediator of the inflammatory response, was mainly secreted by myeloid cells, including macrophages and neutrophils, according to the scRNA-seq data (Fig. 2f). In addition, the expressions of *CCL19/CCL21* and its receptor *CCR7* were most abundant in LNs, when compared with three other regional areas, supporting the key role of the CCL19/CCL21-CCR7 axis in directing lymphocyte homing and in organizing immunological and inflammatory responses in LNs (Extended Data Fig. 3c). Furthermore, expression of co-stimulatory molecules related genes (*CD28* and *CD80*)^36^, which are the most important molecules in T cell expansion, survival, and activation, and inhibitory molecules related genes (*CTLA4*, *BTLA*, *TIGIT*, *CD96*, *CD274*, and *LGALS9*)^36, 37^ vital in T cell inactivation, were both accumulated specifically in LNs, indicating a bidirectional and balanced immune regulation in LNs (Extended Data Fig. 3d, e).

**Fig. 3.**
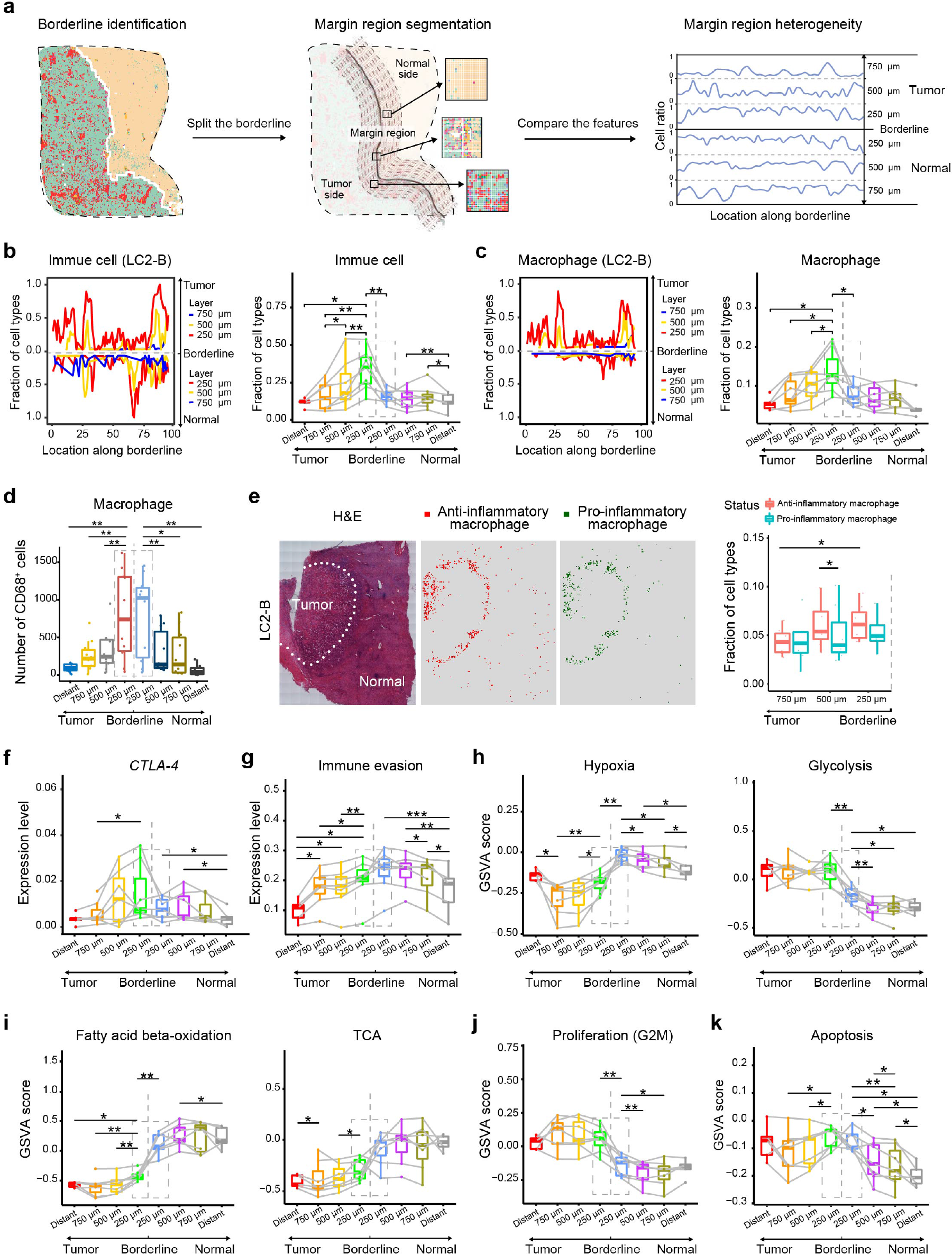
Segmentation of margin areas and distinct features of immune cells and tumor cells at invasive fronts. **(a)** Schematic diagram of the identification of borderlines and segmentation of margin areas to characterize the spatial heterogeneity of margin areas. Three layers with a width of 250 µm were drawn from the normal side and from the tumor side of the borderline, respectively, and each layer was divided into 100 subregions along the borderline. **(b)** Line graphs (left panel) showing the fractions of immune cells in all cell components along the borderlines (100 subregions for each layer) in three layers from the normal side and tumor side of the borderlines (LC2-B), and box plots (right panel) analyzing average fractions of immune cells in all cell components in each layer from seven patients with ICC. “Distant” is defined as a 250 µm-wide zone in tumor tissues or adjacent normal tissues at least 2 mm from the borderline. Two-tailed paired t-tests were used. And only 5 margins included the “distant” area in margin are slides due to the sampling. **(c)** Line graphs (left panel) showing the fractions of macrophages in all cell components along the borderlines (100 subregions for each layer) in three layers from the normal and tumor sides of the borderline (LC2-B), and box plots (right panel) analyzing average fractions of macrophages in all cell components in each layer from seven patients with ICC. Two-tailed paired t-tests were used. (d) Box plots analyzing average number of macrophages (CD68^+^ cells) by immunofluorescence (IF) staining in different layers (1,000 µm in axial length) from the normal and tumor sides of the borderline in 10 ICC patients from Validation Cohort 1. **(e)** Left panel: H&E staining and the corresponding spatial distribution maps of pro-inflammatory and anti-inflammatory macrophages (LC2-B). Right panel: Box plots showing pro-inflammatory and anti-inflammatory macrophage ratios in immune cells in different layers from the tumor side (*n* = 7). **(f)** Box plots showing the expression levels of *CTLA-4* of in different layers of margin areas (*n* = 7). **(g)** Box plots showing the expression levels of immune evasion signatures in malignant cell (tumor side) or all cell components (paratumor side) from different layers of margin areas (*n* = 7). Two-tailed paired t-tests were used. **(h-k)** Box plots showing the expression levels using the Gene Set Variation Analysis (GSVA) score of hypoxia and glycolysis (**h**), fatty acid beta-oxidation and the tricarboxylic acid (TCA) cycle (**i**), proliferative capacity (G2M) (**j**), apoptosis (**k**) of tumor cells in the tumor side and all cell components in the paratumor side of the borderlines (*n* = 7). Two-tailed paired t-tests were used for panel (**b-k**).

### Enrichment of distinctive immune cells, the suppressive immune microenvironment, and metabolic reprogramming of tumor cells at invasive fronts

The area around the borderline, where tumor cells invade paranormal tissues and come into direct contact with a diverse array of stromal cells and ECM components, has been recognized as a complex ecosystem and the most important region of solid tumors for understanding tumor progression, immune surveillance, and infringement of normal tissues^4, 7–11^. The transcriptional landscape in cell and ECM components in four different regional areas also indicated enrichment of inflammatory chemokines/cytokines in the margin areas (Fig. 1b, 2e, and Extended Data Fig. 3a). We observed accumulation of macrophages and NK/T cells close to the boundaries, where tumor cells were in direct contact with adjacent liver tissues, comprising a sophisticated border microenvironment (Extended Data Fig. 4a). Thus, to further characterize the microenvironment at invasive tumor cell fronts, we constructed a model of the precise segmentation of margin areas around the invasive tumor fronts using six layers (each layer was a zone with a width of 250 µm from the borderline in the lateral direction, and each layer was equally divided into 100 equal parts along the axial direction of the borderline) for the bilateral sides (Fig. 3a, see Methods). We first fitted the invasion borderlines of margin areas according to the spatial distribution of tumor cells and hepatocytes in ST slides, which was confirmed by H&E staining of corresponding adjacent slides, and then analyzed the fractions and features of cell components in six layers from the normal and tumor slides of the borderline (Fig. 3a, see Methods). We found that immune cells were recruited to the areas around the borderline from the tumor side, with high heterogeneities in its axial directions, which were especially enriched in the 250 µm wide zone close to the borderline, revealing the high heterogeneities of immune cell distributions around the borderline in both the axial and lateral directions (Fig. 3b). In addition, fibroblasts were found to be enriched in tumor sides, but showed no obvious change in the fractions among different layers of the tumor side (Extended Data Fig. 4b). Among immune cells, macrophages, DCs, T/NK cells, and B cells were found to be more abundant in the areas close to borderlines from the tumor sides (Fig. 3c and Extended Data Fig. 4c). Moreover, the abundance of macrophages was found to be increased from the third layer (750-500 µm) and the second layer (500-250 µm) to the first layer (250-0 µm) in the tumor sides of the borderlines, while there was no change in adjacent normal tissues, further validated by IF result from 10 ICC patients from Validation Cohort 1 (Fig. 3c, d). A significantly increased percentage of anti-inflammatory macrophages (M2-like) rather than pro-inflammatory macrophages (M1-like) was observed from the outer two layers (750–500 µm and 500–250 µm) to the first layer (250–0 µm) close to the borderlines from the tumor sides (Fig. 3e). The percentages of DCs and T/NK cells increased from the third layer (750–500 µm) to the first layer (250–0 µm) close to the borderlines from the tumor sides, while the percentages of B cells increased from the second layer to the first layer (Extended Data Fig. 4c). T cell subtypes, including cytotoxic and exhausted T cells, were scattered among the margin areas without any accumulation around the borderlines (Extended Data Fig. 4d). However, the expression of immune checkpoint genes including *BTLA*, *CTLA4*, *CD96*, and *IDO1* were enriched from the tumor side of the borderlines, indicating the suppressive immune states of immune cells in the area close to the borderline from the tumor side (Fig. 3f and Extended Data Fig. 4e, see Methods). Correspondingly, tumor cells in the first layer showed enhanced immune evasion signatures, when compared with those in outer layers on the tumor side (Fig. 3g). Together, these results showed distinct TME features with enrichment of immune cells, including macrophages (M2-like), T/NK cells, DCs cells, and B cells in the first layer (250–0 µm) from the tumor side as well as a more immune suppressive microenvironment in the closest layers in bilateral sides to the borderline of invasive fronts.

**Fig. 4.**
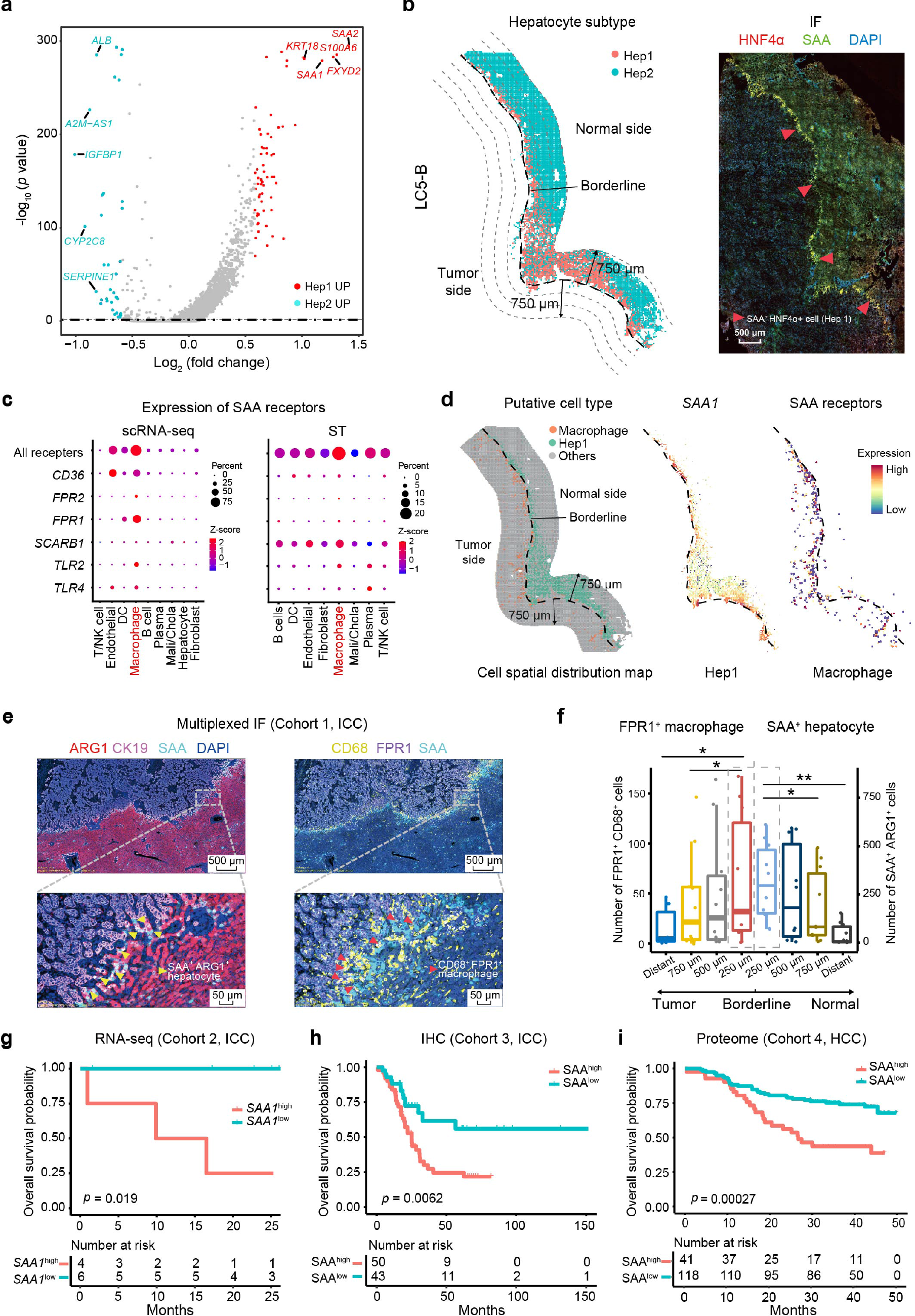
Damaged hepatocyte states at invasive fronts associated with a poor prognosis. **(a)** Volcano plot showing differentially-expressed genes between Hep1 and Hep2. The red dots represent genes upregulated in Hep1 cells and the blue dots represent genes upregulated in Hep2 cells. **(b)** Dot diagram showing the spatial distribution of Hep1 and Hep2 cells in margin areas and Hep1 cells (SAA^high^ hepatocytes) stained by IF staining (HNF4α, SAA, and 4′-6′-diamidino-2- phenylindole (DAPI)) (LC5-B). **(c)** Bubble chart showing expression levels of serum amyloid A (SAA) receptors encoding genes including *CD36, FPR1/2, SCARB1, TLR2*, and *TLR4* in the main cell types in scRNA data and spatially-resolved spatial transcriptomics spots (LC2-B). **(d)** Scatter diagram of the distribution of Hep1 cells (blue dots) and macrophages (red dots) at the LC5-B ST side (left panel). Scatter diagram showing the expression of SAA in Hep1 (middle panel), and the expression of SAA receptors in macrophages (right panel) at the LC5-B ST side. The dash lines represent the borderline. **(e)** Multiplexed IF staining (ARG1, CK19, FPR1, CD68, SAA, and DAPI) showing co-aggregation of FPR1^high^ macrophages (FPR1^+^CD68^+^ cells) and SAA^high^ hepatocytes (SAA^+^ARG1^+^ cells) at invasive fronts of 10 ICC patients from Validation Cohort 1. **(f)** Cell number of macrophages (CD68^+^ cells) and the SAA^+^ hepatocytes (SAA^+^ARG1^+^ cells) in different layers (1000 µm in axial length) at margin areas and distant sites of 10 ICC patients using multiplexed IF staining. **(g)** Overall survival (OS) curves of 10 patients with ICC from Validation Cohort 2, grouped by SAA1 expressions in the border areas (1 cm-wide zone centered on the tumor-normal borderline) using bulk-RNA sequencing. **(h)** OS curves of 93 patients with ICC from Validation Cohort 3 grouped by SAA expressions at invasive fronts using IHC staining. **(i)** OS curves of 159 patients with HCC from Validation Cohort 4, grouped by relative expressions of SAA in paratumor tissues by proteome data.

In addition to the spatial heterogeneities of immune cell distributions and distinct immune suppressive microenvironments in invasive fronts, we further characterized metabolic changes and proliferative capacity of cell components (mainly consisting of hepatocytes) around the borderlines. Generally, tumor cells on the tumor side exhibited lower hypoxia-related pathways and higher glycolysis levels, when compared with all cells in the paratumor side of the borderline (Fig. 3h). The closest bilateral layers to the borderline showed higher hypoxia-related pathway activation, when compared with the corresponding outer regions (Fig. 3h). Consistent with the trend of hypoxia levels, cell glycolysis levels in the first layer on the paratumor side of the borderlines were significantly higher than that in the outer layers (Fig. 3h). Although there was no difference in glycolysis levels of tumor cells among different layers, upregulated levels of tricarboxylic acid cycle and fatty acid metabolism components, including fatty acyl CoA synthesis and fatty acid beta-oxidation of tumor cells, was found in the first layer, when compared with those in outer areas from the tumor side (Fig. 3i and Extended Data Fig. 4f, g). This probably indicated that tumor cells in the first layer underwent metabolic reprogramming to upregulated lipid metabolism because of additional energy requirements for further invasion. As expected, tumor cells on the tumor side of the borderlines exhibited higher G2M scores, when compared with cells (mainly consisting of hepatocytes) at the paratumor side, revealing the higher proliferative capacity of tumor cells (Fig. 3j, see Methods). Apart from a higher proliferative capacity, higher cell apoptosis levels were also detected among the closest layers, when compared with the outer regions from the paratumor side of the borderline, which may have reflected much damaged states of hepatocytes close to the borderline from the paratumor side. For tumor cells, higher apoptosis levels in the closest layer, but no difference of proliferative capacity in different layers from the tumor side of invasive fronts, were observed (Fig. 3j, k). Higher levels of hypoxia, glycolysis, proliferation capacity, and apoptosis in cells (mainly consisting of hepatocytes) were also found in the first layer from the paratumor side. We therefore defined the 500 µm wide zone centered on the borderline as “invasive front,” which was more reasonable than previously defining the 1,000 µm wide area centered on the borderline^7, 9^. Taken together, these results showed an enhanced energy supply from fatty acid metabolism in tumor cells, and a dominant suppressive immune environment was found in the area close to the borderline, especially in invasive fronts.

### The damaged states of hepatocytes at invasive fronts were related to the prognoses of patients with primary and metastatic liver cancers

Apart from the phenomenon of metabolic reprogramming and enhanced immune-escape signatures of tumor cells at invasive fronts, cell components (mainly consisting of hepatocytes) at invasive fronts also showed both enhanced proliferative capacity and increased apoptosis. Some studies have reported that inflammatory responses of hepatocytes contributed to liver cancer or liver metastasis. To characterize the inflammatory response of cell components and to identify distinct cell clusters during tumor invasion, we re-clustered hepatocytes according to their gene expression profiling in the three layers (0–750 µm wide zones) at the paratumor side of invasive fronts. We identified two hepatocyte subtypes (Hep1 and Hep2) according to their different gene expression profiling, where Hep1 cells have remarkablely higher expression levels of *SAA1* (acute phase protein Serum Amyloid A1) and SAA2 (Serum Amyloid A2) compared with Hep2 cells (Fig. 4a). Regarding their spatial distributions, Hep1 cells mainly accumulated in the first layer, with a width of 250 µm close to the borderline, revealing the severely damaged status of hepatocytes because of their direct exposure to tumor invasion, which was confirmed using IF (Fig. 4b and Extended Data Fig. 5a). The bulk RNA-seq data from Validation Cohort 2 also showed much higher expression levels of *SAA1* and *SAA2* around the border areas (bilateral sampling of tissues with a width of 5 mm along the borderline) than corresponding tumor tissues and adjacent normal tissues (n = 10, Extended Data Fig. 5b). Furthermore, IHC results from an additional 93 ICC patients (Validation Cohort 3) also showed higher SAA expressions at invasive fronts than those in outer areas of tumor tissues or adjacent normal tissues (Extended Data Fig. 5c). Using gene enrichment analyses, enriched pathways including oxidative phosphorylation, MYC targets V1, and the epithelial-mesenchymal transition (EMT) for upregulated genes were also identified in Hep1 cells, when compared with Hep2 cells (Extended Data Fig. 5d). Single-Cell Regulatory Network Inference and Clustering (SCENIC) analyses *(45)* also identified distinct upregulated transcriptional factors (TFs) including CEBPG, YY1, ETV6, CDX1, and CTCFL in Hep1 cells, when compared with Hep2 cells (Extended Data Fig. 5e, see Methods). Among these TFs, ETV6, CEBPG, YY1, and BHLHE40, were annotated as potential TFs for *SAA1* and *SAA2* in SCENIC^38^, showing an increased activity score in the first layer close to invasive fronts, which were probably involved in mediating the high expression of *SAA1* and *SAA2* in Hep1 cells (Extended Data Fig. 5f). On the other hand, the reported receptors genes for SAA, including *FPR1*, *FPR2*, *TLR4*, *TLR2*, *SCARB1*, and *CD36*, were predominantly enriched in macrophages according to scRNA-seq data, which was also confirmed by the ST slide results (Fig. 4c). Additionally, both macrophage spots expressing genes encoding SAA receptors and Hep1 cells were spatially accumulated at invasive fronts, further suggesting the recruitment of macrophages for the secretion of SAA from hepatocytes at invasive fronts (Fig. 4d). The accumulation of FPR1^+^ macrophages close to SAA^+^ hepatocytes at invasive fronts was further confirmed by multiplexed IF in Validation Cohort 1, including ICC (n = 10), HCC (n = 5), liver metastasis of colorectal cancer (n = 3), liver metastasis of pancreatic cancer (n = 2), and liver metastasis of lung cancer (n=1) (Fig. 4e, Extended Data Fig. S6 a∼e). Specifically, significantly higher numbers of FPR1^+^ macrophages at the tumor side, and the SAA^+^ hepatocytes at the paratumor side of invasive fronts were found in primary liver cancers including ICC and HCC as well as in metastatic liver cancers mentioned above (Fig. 4f and Extended Data Fig. 6f). Strong correlations between enrichment of macrophages and SAA^+^ hepatocytes at invasive fronts were also observed (R = 0.73, *P = 0.017*) (Extended Data Fig. 6a). In addition, there is also high heterogeneity of invasive fronts along the normal-tumor borderline. We found that tumor cells close to SAA-enriched regions showed an enhanced EMT, upregulated energy metabolic processes (ATP metabolic processes, oxidative phosphorylation, and the respiratory electron transport chain), and activated immune responses (an interferon α response and IL2-STAT5 signaling) compared with the nearby tumor cell Hep1 non-enriched (Hep2 enriched) region at invasive fronts (Extended Data Fig. 7a). Finally, the damaged states of hepatocytes at invasive fronts were found to be related to the prognoses of patients with primary liver cancers. The higher expressions of *SAA1* and *SAA2* in the border areas from Validation Cohort 2 were significantly associated with worse overall survival (OS) of ICC patients (*P* = 0.019 for *SAA1* and *P* = 0.019 for *SAA2*) (Fig. 4g and Extended Data Fig. 7b). IHC results from 93 ICC patients (Validation Cohort 3) also showed significant negative correlations between SAA expression levels at invasive fronts and the OS, as well as relapse-free survival (RFS) for ICC patients (*P* = 0.0062 for OS and *P* = 0.0024; Fig. 4h and Extended Data Fig. 7c). In addition, there was a strong correlation between SAA expressions in invasive fronts and adjacent normal tissues, showing that damaged states of hepatocytes in adjacent tumor tissues were partly reflected at invasive fronts (*R* = 0.75, *P* < 0.001; Extended Data Fig.7d). The higher expression levels of SAA in paratumor areas from Validation cohort 3 were also significantly correlated with shorter OS and RFS in ICC (*P* = 0.0038 for OS and *P* = 0.030 for RFS; Extended Data Fig. 7e, f). The prognostic value of SAA expressions at the transcriptional and protein levels in paratumor tissues of HCC patients was further confirmed by Validation Cohort 4 [RNA-seq data and proteomics data from 159 patients with HBV-related HCC, from our center^28^] (OS, *P* < 0.0001 for *SAA1* or *SAA2* in the RNA-seq data; *P* = 0.00027 for SAA using proteomics analysis) (Fig. 4i and Extended Data Fig. 7g). These results suggested that the damaged states of hepatocytes were associated with tumor progression as well as the prognoses of patients. Most importantly, we identified the damaged states of hepatocytes, which were characterized by overexpression of SAA in response to tumor invasion, followed by recruitment of macrophages for further tumor invasion, ultimately resulting in a worse prognosis.

## Discussion

In this study, we integrated spatially-resolved transcriptomics of subcellular resolution with scRNA-seq to map the transcriptional architecture of four regional sites from eight ICC patients. Compared with other ST methods, our Stereo-seq data had unprecedented nanoscale resolution (220 nm × 220 nm/spot) and expandable areas (10 mm ×10 mm), enabling a more accurate study of cell types or cell states *in situ*, and more space for spatial segmentation and further analyses^12, 20, 39–42^. The use of Stereo-seq in our study enabled the identification of cell types and ECM component distributions in different regions of ICC patients. The results confirmed the compositional heterogeneities of cells and the ECM, as well as their spatial distributions among different regions. By constructing a segmentation model of areas around the tumor margin area, we firstly identified the distinctive enrichment of immune cell clusters, metabolic reprogramming of tumor cells, and inflammatory states of hepatocytes in the areas around invasive fronts (500 µm wide zone centered on the borderline, 250 µm each side).

Invasive fronts, where tumor cells invade into paratumor tissues and come into direct contact, have been recognized as the most important regions of solid tumors necessary for understanding tumor progression and metastasis^4, 7–11, 20^. Comprehensive mapping of the transcriptional architecture of these areas has facilitated a better understanding of the molecular pathology, and development of personalized therapeutic strategies, such as immune checkpoint blockade therapy for solid tumors. As a fundamental biological feature of the TME, spatial heterogeneity has been largely ignored due to the limitations of research tools^5, 6, 12, 43, 44^. In the present study, based on comprehensive mapping of the spatial transcriptomics architecture using Stereo-seq, the spatial heterogeneities of invasive fronts were compared with other regions in ICC, which helped to understand tumor behaviors, including invasion and metastasis. Based on a constructed model consisting of three layers on bilateral sides of the borderlines in ST slides, we found that enrichment of immune cells and a distinctive immune suppressive microenvironment and metabolic reprogramming of tumor cells were observed in the closest 250 µm wide layer from the tumor side of the borderlines. Among the immune cells, macrophages mainly comprised of the dominant M2-like macrophage accumulated at invasive fronts, indicating the dominant promotion of tumor progression by anti-inflammatory macrophages. Furthermore, we found enhanced expressions of immune checkpoint genes, including *CTLA4* in immune cells, and we also found upregulated immune evasion signatures in tumor cells in the closest layer to the borderlines. Thus, a more suppressive immune microenvironment in invasive fronts was found, which could facilitate further tumor invasion and metastasis of solid tumors.

Based on precise segmentation of margin areas, our study also comprehensively characterized the distinct cell states of tumor cells at invasive fronts. By analyzing conventional pathways related with tumor cell behaviors, we observed more energy supply components from upregulated fatty acid metabolism, including increased fatty acid synthesis and oxidation in tumor cells in the closest layer at the tumor side, when compared with outer layers from the tumor-paratumor borderline, further indicating that tumor cells at these locations underwent a distinct metabolic reprogramming of enhanced fatty acid metabolism for an increased energy supply, to better facilitate tumor cell invasion. Meanwhile, we also found enhanced hypoxia states, glycolysis levels, and both proliferative capacity and apoptosis levels in cell components at the closest layer from the paratumor side of the borderlines, when compared with outer areas. Because all of these results revealed distinct inflammatory states and special metabolic features at the areas around invasive fronts, we redefined the 500 µm wide zones centered at the tumor borderline as “invasive fronts,” which was more reasonable than previously defining a 1,000 µm wide area centered on the borderline^7–9, 20^. Furthermore, metabolic reprogramming of enhanced fatty acid metabolism in tumor cells among invasive fronts for an increased energy supply could account for further tumor invasion of solid tumors. Therefore, therapeutic approaches that blocked this process could be a novel treatment to inhibit tumor invasion and metastasis.

By further characterizing features of cell components at invasive fronts, we found that hepatocytes at invasive fronts also underwent distinct inflammatory changes, with high expressions of SAA and their TFs, including ETV6, CEBPG, YY1, and BHLHE40, which reflected the damaged conditions of hepatocytes invaded by tumor cells^45, 46^. SAA can directly coordinate accumulation of innate immune cells like monocytes, immature DCs, neutrophils, and T cells. Receptors for SAA, including FPR1 and FPR2, were detected in macrophages at invasive fronts, and significantly accumulated close to Hep1 cells, which probably resulted from the recruitment of macrophages close to the borderline because of secretion of SAA by damaged hepatocytes. This possibility is consistent with a study reporting that hepatocytes upregulated SAA1 expression and provided a pro-metastatic niche for liver metastasis of pancreatic cancer in a mouse model^45^. The high expression and enrichment of SAA in invasive fronts could explain the aggregation of immune cells at this site, which has been reported in other studies, as well as our own study. SAA has been reported to contribute to M2-macrophage differentiation^47^, which could account for the presence of more M2-like macrophages found at invasive fronts. Moreover, higher expressions of SAA and accumulation of macrophages were also confirmed at invasive fronts in other solid tumors including HCC, liver metastasis of colorectal cancer, and pancreatic cancer, and was further found to be related with worse prognoses of patients. Our work identified heterogeneity of invasive fronts in the axial direction, with high variations of tumor cell states and damaged hepatocyte distributions along the borderlines. Most importantly, we uncovered a novel role of SAA expressions for hepatocytes at invasive fronts, which reflected the damaged states of hepatocytes, and mediated recruitment of macrophages (mainly M2-like macrophages) to further promote tumor progression.

Although Stero-seq provided a high nanoscale resolution (220 nm × 220 nm/spot) to capture transcripts of a few hundred spots per cell, it was still difficult to identify cell boundaries and capture exact transcripts at the level of single cells^21, 38, 39^. In our study, the raw spatial expression matrix at a nanoscale resolution was converted into cell-sized pseudo-spots of 25 µm square sizes (50 × 50 bins/pseudo-spot) and assigned to specific cell types by the highest probability. Thus, it was still difficult to spatially map resolved single cell genetic information and precisely infer cell types because of technical limitations.

In conclusion, this study characterized the complexity and heterogeneity of tumor ecosystems, as well as cellular interactions in different regional sites of ICC, which identified immune cell infiltration, a suppressive immune microenvironment, metabolic reprogramming of tumor cells, and damaged hepatocytes enriched at invasive fronts. We also verified this findings in HCC, liver metastasis of colorectal or pancreatic cancer. Overall, the results provided a comprehensive understanding and detailed characterization of the heterogeneity present at tumor invasive fronts, which will help to develop more precise and effective therapeutic targets for the treatment of solid tumors.

## Acknowledgments

We thank X. Tong, M. Zhan and W. Chen for helpful advice and discussions. We also acknowledge members of the National Center for Protein Science Shanghai for assistance in microscopy.

## Funding

This work was jointly supported by the National Key Research & Development Program of China (2019YFC1315800 and 2019YFC1315802), the State Key Program of National Natural Science of China grants (81830102), the National Natural Science Foundation of China grants (81772578, 81772551, 81872355, 81830102, 81802991, 82150004, and 82072715), the Shanghai Municipal Health Commission Collaborative Innovation Cluster Project (2019CXJQ02), the Shanghai “Rising Stars of Medical Talent” Youth Development Program (Outstanding Youth Medical Talents), the projects from the Shanghai Science and Technology Commission (19441905000 and 21140900300), and Shenzhen Key Laboratory of Single-Cell Omics (ZDSYS20190902093613831).

## Author contributions

Each author’s contribution to the paper was listed below.

Conceptualization: JZ, XRY, AC, LQL, JF

Data curation: XXZ, ZHJ

Formal analysis: XXZ, ZHJ, LZH, JYY, YZ, XRZ, MYZ

Resources: FYC, JYY, JZ, XRY, AC, AH, SPL, LQL, XX, JW, HXS, SL, YXL, XDW, SY

Software: XXZ, YQB, LZH

Validation: JYY, YFC, XXZ, YZ, YJ, RKL, QQL, AH, DZG, PXW, HXS, ZFJ, NY, YY, YL, FML

Methodology: YQB, JSX, MNC, AC Investigation: LW, JSX, JYY, FYC

Visualization: XXZ, ZHJ, YZ, YQB, JYY, JSX, LW, WDH, HY, JJL

Funding acquisition: AC, JZ, SPL, XRY, XX, JW

Project administration: LW, JYY, LQL, XRY

Supervision: JZ, AC, XRY, SPL, JF, LQL, XX, JW, HMY

Writing – original draft: LW, JYY, YQB, FYC, XXZ, JSX, LZH

Writing – review & editing: LW, JYY, XRY, YZ, YQB, XXZ, JSX, SPL

## Competing interests

Authors declare that they have no competing interests.

## Data and materials availability

The ST data and scRNA-seq data generated by this study will be available in CNGB Sequence Archive (CNSA). The bulk RNA-seq and proteome data of Validation cohort 4 ^28^ are available with accession number OEP000321 in The National Genomics Data Center (NGDC).

## Materials and Methods

### Study subjects

Matched, fresh tumor tissues, adjacent normal tissues, margin area tissues, and lymph node samples were collected as the **Discovery Cohort** from 8 ICC patients for spatial transcriptomics analyses, with detailed and pathological information shown in Supplementary Table 1. Formalin-fixed paraffin-embedded (FFPE) tissue blocks of margin area tissues were collected from twenty-three patients who had undergone liver resection and were pathologically diagnosed with liver cancer (ICC, *n* = 10; HCC, *n* = 5; liver metastasis of colorectal cancer, *n* = 3; liver metastasis of pancreatic cancer, n = 2, and liver metastasis of lung cancer, *n* = 1) as **Validation Cohort 1**. Matched, frozen tumor tissues (at least 2 cm from the tumor-normal borderline), adjacent normal tissues (at least 2 cm from the tumor-normal borderline), and border area tissues (1 cm-wide zones centered on the tumor-normal borderline) samples were collected from 10 ICC patients who had undergone liver resection with pathological diagnoses of ICC as **Validation Cohort 2**. FFPE tissue blocks of margin area tissues (2 × 2 × 1 cm, 2 cm-wide zones centered on the tumor-normal borderlines) samples were collected from 93 ICC patients who had undergone liver resection with pathological diagnoses of ICC as **Validation Cohort 3**. Paired frozen tumor tissues and paratumor tissues were collected from 159 HCC patients who had undergone liver resection with pathological diagnoses of HCC as **Validation Cohort 4** ^28^. The available clinical features of Validation cohorts are summarized in Supplementary Table 3. All patients provided informed consent for collection of clinical information, and the tissue collection protocols were approved by the Institutional Review Board [approval B2018-018(2)] at Zhongshan Hospital Fudan University.

### Single-cell RNA sequencing

#### Single-cell isolation

Margin area tissues (a 1 cm-wide zone centered on the tumor-normal borderline) were surgically removed from resected liver lobes from ICC patients and immersed in a complete medium containing 90% Dulbecco’s Modified Eagle medium (Gibco, Gaithersburg, MD, USA) and 10% fetal bovine serum (FBS; Gibco), and transported to the laboratory in a refrigerated container. Suitable small tissue blocks were then cut into pieces, which were transferred to MACS C tubes (Miltenyi Biotec, Bergisch Gladbach, Germany), with 5 mL of digesting enzyme included in a Tumor Dissociation Kit (Miltenyi Biotec). The tissues were then converted into single-cell suspensions using a gentle MACS Dissociator (Miltenyi Biotec) with the following steps: milled; incubated at 37°C for 30 min on a shaker; milled; incubated at 37°C for 30 min; milled; filtered through a 70 mm filter in 2% FBS. Finally, the single-cell suspension was centrifuged at 300 × g for 7 min, and resuspended with cell resuspension buffer at a cell concentration of 1,000 viable cells/µl.

#### Single-cell RNA-seq library construction

Single-cell RNA-seq libraries were prepared using DNBelab C4 system as previously described^39^. Barcoded mRNA capture beads, droplet generation oil, and the single-cell suspension were loaded into the corresponding reservoirs on the chip for droplet generation. The droplets were gently removed from the collection vial and placed at room temperature for 20 minutes. Droplets were then broken and collected by the bead filter. The supernatant was removed, and the bead pellet was resuspended with 100 µl RT mix. The mixture was then thermal cycled as follows: 42 °C for 90 minutes, 10 cycles of 50 °C for 2 minutes, 42 °C for 2 minutes. The bead pellet was then resuspended in 200 µl of exonuclease mix and incubated at 37 °C for 45 minutes. Afterward, the PCR master mix was added to the beads pellet and thermal cycled as follows: 95 °C for 3 minutes, 13 cycles of 98 °C for 20 s, 58 °C for 20 s, 72 °C for 3 minutes, and finally 72 °C for 5 minutes. Amplified cDNA was purified using 60 µl of AMPure XP beads. The cDNA was subsequently fragmented to 400-600 bp with NEBNext dsDNA Fragmentase (New England Biolabs) according to the manufacturer’s protocol. Indexed sequencing libraries were constructed using the reagents in the C4 scRNA-seq kit following the steps: (1) post fragmentation size selection with AMPure XP beads; (2) end repair and A-tailing; (3) adapter ligation; (4) post ligation purification with AMPure XP beads; (5) sample index PCR and size selection with AMPure XP beads. The barcode sequencing libraries were quantified by Qubit (Invitrogen). The sequencing libraries were sequenced using the DIPSEQ T1 sequencer at the China National GeneBank. The read structure was paired-end with Read 1, covering 30 bases inclusive of the 10 bp cell barcode 1, 10 bp cell barcode 2, and 10 bp unique molecular identifier, and Read 2 containing 100 bases of transcript sequence, and a 10 bp sample index.

### Methods of Spatial Transcriptomics

#### Stereo-seq chip preparation

Generation of capture chips was performed following the Stereo-seq protocol ^39^. In brief, to generate the DNB array for *in situ* RNA capture, we first synthesized random 25 nucleotide CID-containing oligonucleotides, circularized with T4 DNA ligase, and splint oligonucleotides. DNB were then generated by rolling circle amplification and loaded onto the patterned chips (65 mm × 65 mm). Next, to determine the distinct DNB-CID sequences at each spatial location, single-end sequencing was performed on a DNBSEQ-Tx21 sequencer (MGI Research) with sequencing SE25 strategy. After sequencing, poly-T and 10 bp MID-containing oligonucleotides were hybridized and ligated to the DNB on the chip. This procedure produced capture probes containing a 25 bp CID barcode, a 10 bp MID, and a 22 bp poly-T ready for *in situ* capture. CID sequences together with their corresponding coordinates for each DNB were determined using a base calling method according to the manufacturer’s instruction for the DNBSEQ sequencer. After sequencing, the capture chip was split into smaller size chips (10 mm × 10 mm). At this stage, all duplicated CID that corresponded to non-adjacent spots were removed.

#### Tissue sectioning, fixation, staining, and imaging

Matched, fresh tumor tissues (2 × 2 × 1 cm, at least 2 cm from the tumor-normal borderline), adjacent normal tissues (2 × 2 × 1 cm, at least 2 cm from the tumor-normal borderline), margin area tissues (2 × 2 × 1 cm, a 2 cm-wide zone centered on the tumor-normal borderline), and LN samples were collected and snap-frozen in optical cutting tissue (OCT) compound (Tissue-Tek; Sakura Finetek USA, Torrance, CA, USA). After collection, the tissues were snap-frozen in liquid nitrogen containing prechilled isopentane in Tissue-Tek OCT and transferred to a -80°C freezer for storage before the experiment. The prefrozen liver tissues in OCT were transverse sectioned at a thickness of 10 µm using a CM1950 cryostat (Leica, Wetzlar, Germany). Tissue sections adhering to the Stereo-seq chip surface were incubated at 37°C for 3 min. The tissues were then fixed in methanol and incubated at -20°C for 40 min. The adjacent tissue sections adhering to the glass slides were stained using H&E. Imaging for both procedures was conducted using a Ti-7 Eclipse microscope (Nikon, Tokyo, Japan).

Tissue patches on the chip were permeabilized using 0.1% pepsin (Sigma-Aldrich, St. Louis, MO, USA) in 0.01 M HCl buffer (pH = 2), incubated at 37°C for 10 min and then washed with 0.1× SSC buffer (Thermo Fisher Scientific) supplemented with 0.05 U/µL RNase inhibitor (New England Biolabs, Ipswich, MA, USA). Released RNA from permeabilized tissues was captured using DNB probes and reverse-transcribed overnight at 42°C using in-house SuperScript II (10 U/µL reverse transcriptase), 1 mM dNTPs, 1 M betaine solution PCR reagent, 7.5 mM MgCl2, 5 mM DTT, 2 U/µL RNase inhibitor, 2.5 µM Stereo-TSO, and 1× First-Strand buffer. After in situ reverse transcription, tissue patches were washed twice with 0.1× SSC buffer and digested with tissue removal buffer (10 mM Tris-HCl, 50 mM EDTA, 200 mM NaCl, and 1.5% SDS) at 37°C for 30 min. The cDNA-containing chips were then incubated with 400 µL cDNA release buffer (10 mM Tris-HCl, 50 mM EDTA, 200 mM NaCl, 1.5% SDS, and 40 U/mL Proteinase K) treatment for 3 h at 55°C, and then washed once with 400 µL of 0.1× SSC buffer. All products were purified using 0.8 × Ampure XP Beads (Vazyme Biotech, Nanjing, China), and were amplified with KAPA HiFi Hotstart Ready Mix (Roche, Basel, Switzerland) using 0.8 µM cDNA-PCR primers. PCR reactions were conducted as follows: first incubation at 95°C for 5 min, 15 cycles at 98°C for 20 s, 58°C for 20 s, 72°C for 3 min, and a final incubation at 72°C for 5 min. PCR products were purified using 0.6 × Ampure XP Beads. The concentrations of cDNA were quantified using a Qubit™ dsDNA Assay Kit (Thermo Fisher Scientific).

#### Library preparation and sequencing

A total of 20 ng of cDNA was fragmented with in-house Tn5 transposase at 55°C for 10 min. The reactions were then terminated by the addition of 0.02% SDS buffer with gentle mixing at 37°C for 5 min. Fragmentation products were amplified as follows: 25 µL of fragmentation product, 1× KAPA HiFi Hotstart Ready Mix, 0.3 µM Stereo-Library-F primer, 0.3 µM Stereo-Library-R primer in a total volume of 100 µL with the addition of nuclease-free H2O. The reaction was then run as follows: one cycle at 95°C for 5 min, 13 cycles at 98°C for 20 s; 58°C for 20 s and 72°C for 30 s), and one cycle at 72°C for 5 min. The PCR products were purified using Ampure XP Beads (Vazyme; 0.6× and 0.2×) for DNB generation and finally sequenced (a paired-end of 100 bp) using a MGI DNBSEQ-Tx sequencer.

#### Cell clustering

Clustering analysis of the scRNA dataset was performed using Seurat (version 3.2.2) and the R program, and the parameters were manually curated to portray an optimal classification of cell types with empirical knowledge. Specifically, low quality cells were filtered with fewer than 500 detected genes or above 6,000, as well as with higher than 20% mitochondrial counts in data preprocessing, and all query genes were guaranteed to be expressed in at least three cells for further use. The top 3,000 highly variable genes were then selected according to their mean-variance ratio on expression levels after log1p normalization. For downstream clustering and visualization, principal component analysis (PCA)-based dimension reduction was initially generated, and the first 18 PCs were extracted for subsequent Louvain clustering to define the cell types (the resolution was set to 0.3). The clustering result was finally characterized in a two-dimensional space using the UMAP technique, and the cell types were annotated by known biomarkers that were more highly expressed in a particular cluster (via FindAllMarkers function with default parameters).

#### Cell type inference of spatial transcriptome spots

To overcome the low RNA capture efficiencies on single DNB spots at a resolution of 500 nm, the raw spatial expression matrix was convoluted into larger pseudo-spots of 50 × 50 window size (bin50 for short), or more precisely, the 25 µm squares. The cell type composition for each bin50 spot was then inferred by SPOTlight software ^31^ (version 0.1.6), with factorized cell type-specific topic profiles from paired scRNA-seq data. The potential composition of each spot was pruned and renormalized using the top four cell types with respective probabilities in descending order, and the primary cell type was assigned for visualization.

#### Differential gene expression analysis

Differentially expressed gene (DEG) analysis in each cluster was performed using the FindAllMarkers function of the Seurat package (v3.2.2), and the differential expression genes between the two groups are detected by FindMarkers function. The parameter condition is min.pct=0.1, logfc.threshold=0.15.

#### Functional enrichment analysis

To identify the biological function of differentially-expressed genes of each cluster, we performed differential expression gene set enrichment analyses using the Molecular Signatures Database of H (hallmark gene sets, version 7.4) by Gene Set Enrichment Analysis ^48^ (GSEA, https://www.gsea-msigdb.org/gsea/index.jsp). To characterize the differences in pathways as well as biological functions between tumor cells from SAA-enriched area and non-SAA-enriched area, the differentially-expressed genes between the tumor cells in two layers around borderline of LC5-B were used. Similarly, the differential genes of the two subtypes of hepatocytes in LC5-B were used for pathway enrichment analysis.

#### Transcription factors analysis

We prepared all bins (bin 50) from stereo-seq. A bin to gene signal matrix depicting gene abundances was input into pySCENIC pipeline^38^ with default settings to infer statistically active TFs and their targets. First, it inferred co-expression modules using GRNBoost2, a regression per-target approach. Second, it pruned indirect targets from modules using regulatory motif discovery (cisTarget). In brief, enriched motifs were discovered from all genes in co-expression modules. Each remaining TF and its potential direct target was called regulon. Finally, it used AUCell as a metrics computing the activity of each regulon in each bin. To obtain each TF’s activity scores, we put the bin matrix into pySCENIC pipeline^38^. To identify the specific regulons of hepatocyte subtypes, we used Wilcoxson Rank-Sum test to calculated the different TF between Hep1 and Hep2.The specific TF in Hep1 were displayed by Feature Plots on spots.

### Survival analysis

The samples of patients for survival analyses were split into two groups (high and low) according to the quantile expressions of the proposed gene(s) in the surv_cutpoint function of the R survminer package. Kaplan-Meier survival curves measuring the fractions of patients living for a certain time were plotted to compare the two patient cohorts and assess the effect of the particular gene(s) on prognoses. Statistical significance was calculated using the log-rank test. All analyses were performed using the R 3.6.0 framework.

### Identification of the tumor-normal border and invasive fronts

To identify the border of peritumor and intratumor regions by spatial transcriptome profiling, the tumor section was first processed into a binary image that masked the predicted hepatocyte cells. Only the large area of the normal liver tissue was retained as a region of interest, and scatter signals were filtered by window-size pixel thresholding (the threshold was set manually to sweep scatter signals outside regions on different sections), and the boundary pixels were extracted using the Contours function in the Python OpenCV package and initialized into a rough edge. The edge was then smoothed by a spline fitting (with 20 degrees of freedom) using the R spline package, and local segmentation and/or rotations were introduced on the complex borders, which could not be directly fitted (for example, a border graphed as a parabola with a horizontal axis of symmetry was barely fitted without a 90° rotation).

After the determination of the border, parallel curved lines were generated by perpendicularly extending 250, 500, and 750 µm to both tumor and normal sides, to measure the appropriate width of the tumor invasive front. To be specific, six infiltrating layers (bidirectional) were derived from the border, and each layer was segmented into 100 tiles with approximately equal areas along the border line. The spots/cells were subsequently assigned to the corresponding tiles by calculating the sign of the outer product of their centroid coordinates to each edge of the tiles, and were used to assess the variations of spatial gradients of cellular components and gene expression profilers on both tangential and normal directions of invasive front.

### Detection of cell subtypes

#### T cell, B cell, macrophage, and CAF subtypes

To study the subclass of each cell type, the relative higher confidential cell spots were extracted from the SPOTlight output ^31^, under criteria that the normalized probability of primary predicted cell types was ≥ 0.4 and secondary types was ≤ 0.2. CPM normalization was generated for each spot to ensure the gene relative abundance between comparable spots, and gene standardization was applied to the empirical biomarkers of corresponding cell subtypes (summarized in Supplementary Table 4) to balance the expression levels for those selected marker genes that were significantly expressed in corresponding subtypes. The highest average score was then assigned to the spot to represent the most probable subtype for downstream analysis.

#### Identification of SAA1/2^high^ hepatocyte subtypes

The SAA1/SAA2-abundant hepatocytes were directly distinguished by increasing the resolution of Louvain unsupervised clustering in invasive fronts of LC2 and LC5 samples, respectively, and the most significant marker genes (including *SAA1* and *SAA2*), which defined this particular hepatocyte subtype, were intersected into a reference set. In a similar manner, the countered reference set was generated, specifying hepatocyte cells with SAA1/2 not significantly expressed. To identify the *SAA1/2*-abundant hepatocytes on other tumor border sections, the genes in the two reference sets were normalized and standardized, and a summed scaled expression level of two cell clusters was used to define the SAA1/2^high^ or SAA1/2^low^ hepatocytes.

#### Cell type enrichment, gene expression, and tumor hallmark score analysis at invasive fronts

After stratification and blocking of the tumor invasive front into small quasi-equal-area tiles, the cell type composition, gene expression, and tumor hallmark scores were assessed on these elaborated spatial bulk RNA profilers. In particular, cell types were summarized by the normalized probabilities of each bin50 spot that was inferred by the former SPOTlight^31^ results. The gene expressions were normalized using the CPM and compared between the tiles and layers on a bulk level. The tumor hallmark scores were generated by extracting the tumor cell spots (under the criteria of cell subtype detection section) in border-to-intratumor tiles, normalizing, and comparing to border-to-peritumor tiles by Gene Set Variation Analysis (GSVA) analysis. All results were shown as percentages of each tile, and then statistical analyses were conducted to study the heterogeneity on both tangential and normal directions of tumor invasive fronts.

#### Multiplexed immunofluorescence staining

In Validation Cohort 1, FFPE tissue blocks of margin area tissues (2 × 2 × 1 cm, a 2 cm-wide zone centered on the tumor-normal borderline) were collected from 21 patients who had undergone liver resection and were pathologically diagnosed with liver cancer. Multiplex staining of FFPE tissues was conducted using a PANO 7-plex IHC kit (Panovue) according to the manufacturer’s instructions. Primary antibodies were to CK19 (#ab52625; Abcam, Cambridge, UK), ARG1 (#93668S, Abcam), SAA1/2 (#ab207445, Abcam), FPR1 (ab113531, Abcam), CD68 (#76437S, Cell Signaling Technology, Danvers, MA, USA), S100P (#ab124743, Abcam), CK20 (#13063S, Abcam), TTF1 (#ab76013, Abcam) were sequentially used, followed by incubation with horseradish peroxidase-conjugated secondary antibody and tyramide signal amplification. The slides were then microwave heat-treated after each TSA procedure. Nuclei were stained with 4′- 6′-diamidino-2-phenylindole (DAPI; Sigma-Aldrich) after all the human antigens had been labelled. For each slide, three zones with a width of 500 µm and length of 1,000 µm from tumoral areas, paratumor areas, and the areas centered on the borderline were selected for image capture.

#### Bulk RNA extraction and sequencing

For 10 ICC patients who had undergone liver resection and were pathologically diagnosed with ICC in **Validation Cohort 2**, matched, frozen tumor tissues (at least 2 cm from the tumor-normal borderline), adjacent normal tissues (at least 2 cm from the tumor-normal borderline), and border area tissues (1 cm-wide zones centered on the tumor-normal borderline) were collected. Total RNAs of tumor tissues, border area tissues, and adjacent normal tissues from 10 ICC patients were isolated using a RNeasy Mini Kit (Qiagen, Hilden, Germany). Strand-specific libraries were prepared using a TruSeq Stranded Total RNA Sample Preparation kit (Illumina, San Diego, CA, USA) following the manufacturer’s instructions. Briefly, mRNA was enriched with oligo(dT) beads. Following purification, the mRNA was fragmented into small pieces using divalent cations at 94°C for 8 min. The cleaved RNA fragments were then copied into first strand cDNA using reverse transcriptase and random primers, followed by second strand cDNA synthesis using DNA Polymerase I and RNase H. These cDNA fragments then underwent an end repair process, involving addition of a single “A” base, followed by ligation of the adapters. The products were then purified and enriched with PCR to create the final cDNA library. Purified libraries were quantified using a Qubit 2.0 fluorometer (Life Technologies, Carlsbad, CA, USA) and confirmed using a 2100 Bioanalyzer (Agilent Technologies, San Jose, CA, USA) to confirm the insert size and calculate the mole concentration.

Clusters were generated by cBot with the library diluted to 10 pM and then sequenced using a NovaSeq 6000 (Illumina). The library construction and sequencing were performed at Shanghai Biotechnology Corporation (Shanghai, China).For each sample, 33–95 million RNA-seq clean reads were obtained using HISAT2 (hierarchical indexing for the spliced alignment of transcripts)^49^, version 2.0.477. Sequencing read counts were calculated using Stringtie ^50, 51^, version 1.3.0. Expression levels from different samples were then normalized by the Trimmed Mean of M values method. The normalized expression levels of different samples were converted to fragments per kb of transcript per million mapped (FPKM) fragments.

#### Immunohistochemical staining and evaluation

FFPE tissue blocks of margin area tissues (2 × 2 × 1 cm, a 2 cm-wide zone centered on the tumor-normal borderline) were collected from 93 ICC patients who had undergone liver resection and were pathologically diagnosed with ICC as **Validation Cohort 3**. FFPE tissue blocks of margin areas from 93 patients (**Validation Cohort 2**) were used for IHC staining. Primary antibody against SAA (#ab190802, Abcam) was used for the staining of SAA. All staining was conducted using the IHC/ISH System (BenchMark GX; Roche, Basel, Switzerland) following the manufacturer’s instructions.

For the evaluation of the staining index, three zones with a width of 500 µm and a length of 1 mm were selected from tumor areas, areas around the borderline (the zone was centered on the borderline), and paratumor areas. The staining index was further acquired using the ImageJ 1.53 (National Institutes of Health, Bethesda, MD, USA) IHC profiler.

### Statistical analysis

Statistical analyses were performed using the R 3.6.0 framework, including Student’s t-test, Wilcoxon’s sign rank test, and Wilcoxon’s rank-sum test. Asterisks represented the significance levels of the performed tests (\**P* <0.05; \*\**P* < 0.01; \*\*\**P* < 0.001).

---

**Extended Data Fig 1.**
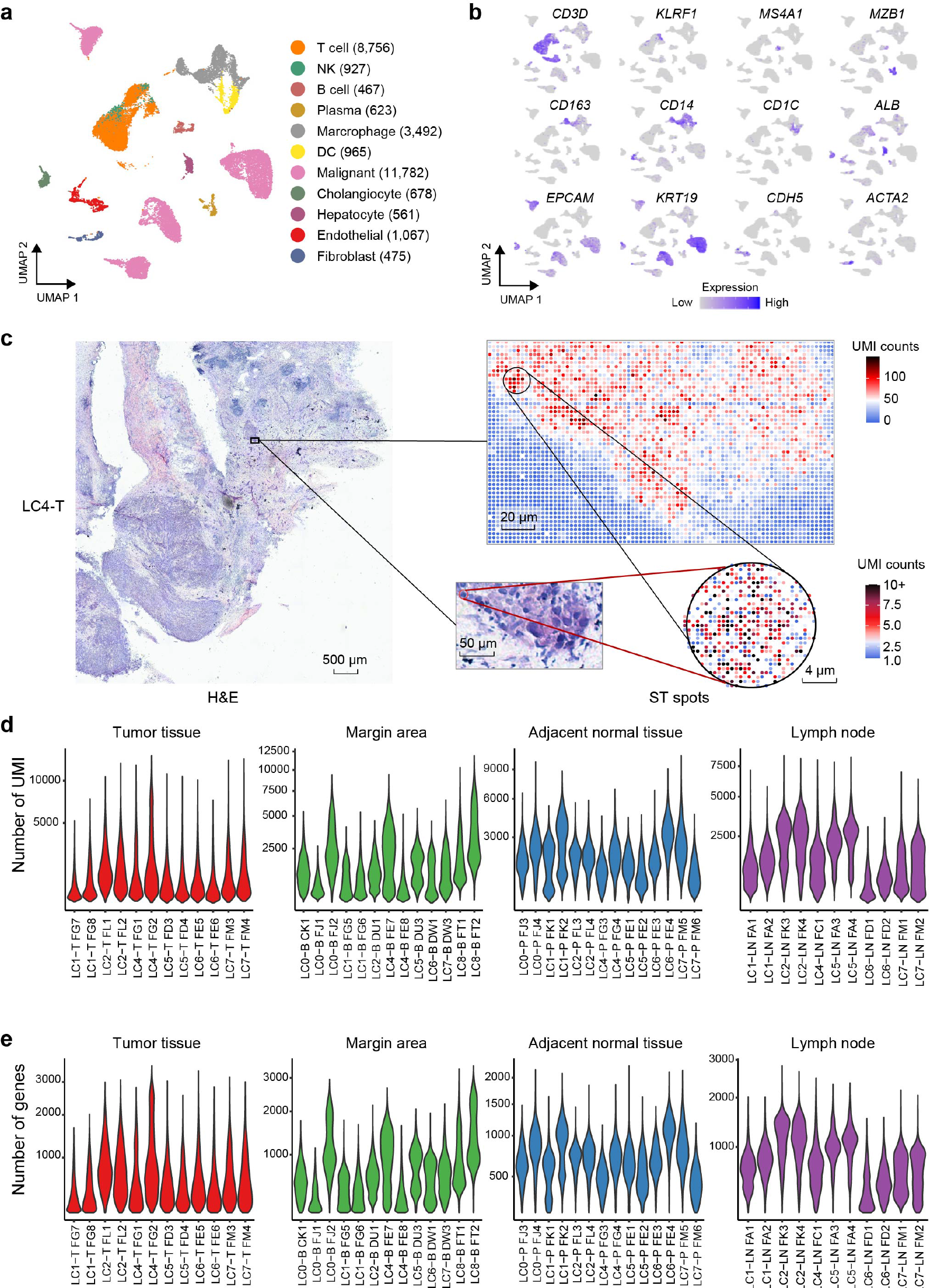
Cell type assignment of scRNA-seq profiling and spatial transcriptome profiling abundance in different intrahepatic cholangiocarcinoma (ICC) regions. (a) An UMAP plot for the cell type identification of 29,189 single cells based on scRNA-seq data using Seurat. **(b)** UMAP plots color-coded for the expression (gray to purple) of marker genes for distinct cell types. **(c)** Spatial transcriptomics (ST) spots were mapped to H&E staining of adjacent slides of LC4-T, and the raw spatial expression matrix with each bin/spot was convoluted into pseudo-spots with 2.5 µm squares (5 × 5 bins/spot) to show unique molecular identifiers (UMI) counts of the cell nucleus (with a diameter of 15 µm) of a tumor cell by ST spots. **(d-e)** Violin plots showing UMI counts (**d**) and numbers of genes (**e**) captured in 50 ST slides of 27 samples (T, *n* = 6; P, *n* = 7; B, *n* = 8; LN, *n* =6) from eight patients with ICC.

**Extended Data Fig 2.**
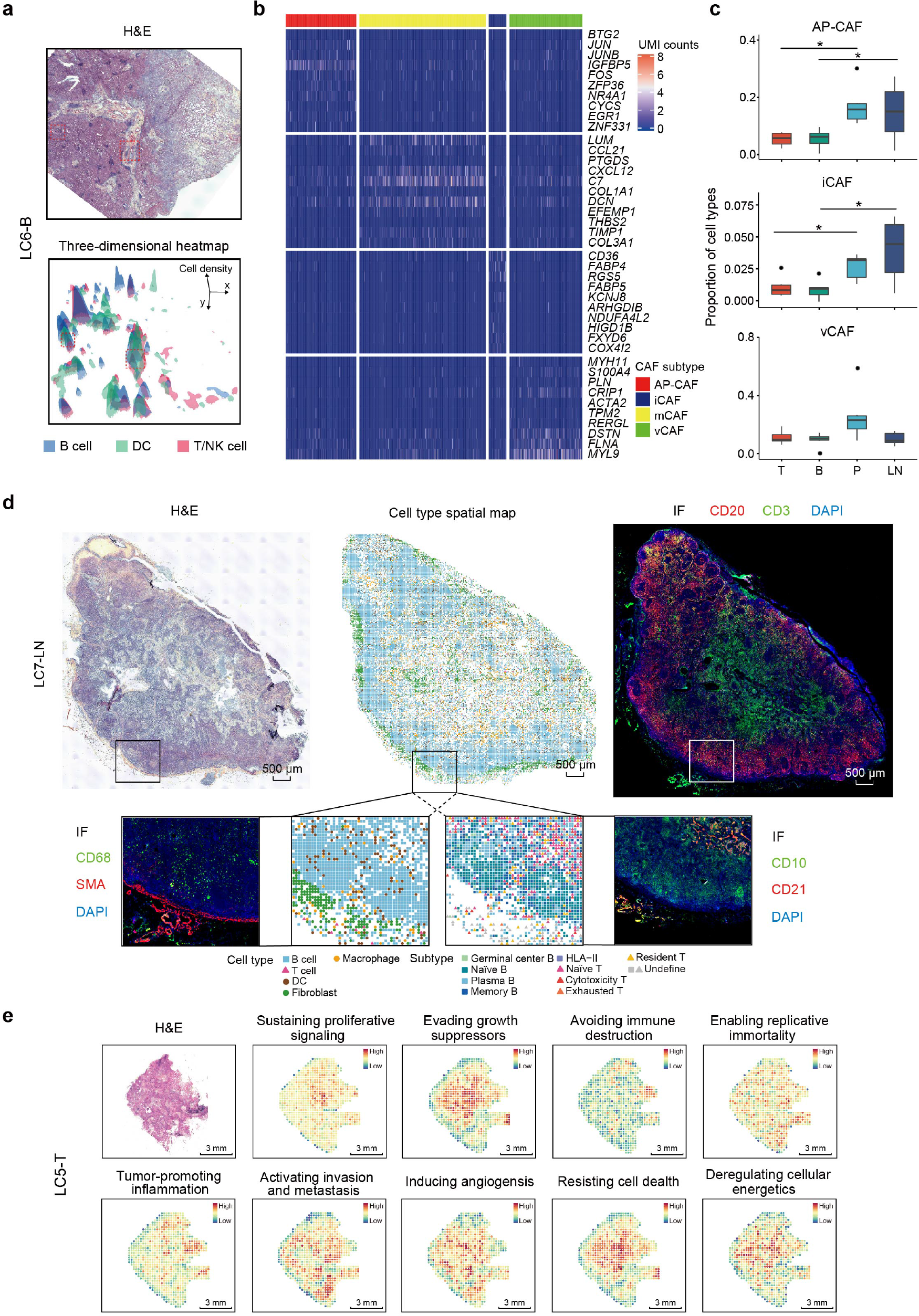
The heterogeneities of spatial distributions of cell components in four regional sites of patients with ICC. **(a)** H&E staining and corresponding heat maps illustrating the location of tertiary lymphoid structures (TLS) featured with co-aggregation of B cells, T/NK cells, and DCs in LC6-B. **(b)** Heat map showing differential expression genes of cancer-associated fibroblast (CAF) subtypes AP-CAF, i-CAF, mCAF, and vCAF in tumor tissues. **(c)** Box plots showing the percentages of AP-CAF, iCAF, and vCAF subtypes in all fibroblast in four regional sites of eight patients with ICC. **(d)** The H&E staining, cell type spatial maps and corresponding IF illustrating the structure of the lymph node including the germinal center and the distribution of subpopulations of main cell types in lymph nodes (LC7-LN). **(e)** H&E staining and signature of cancer hallmarks for corresponding spatial areas of tumor tissue LC5-T.

**Extended Data Fig. 3.**
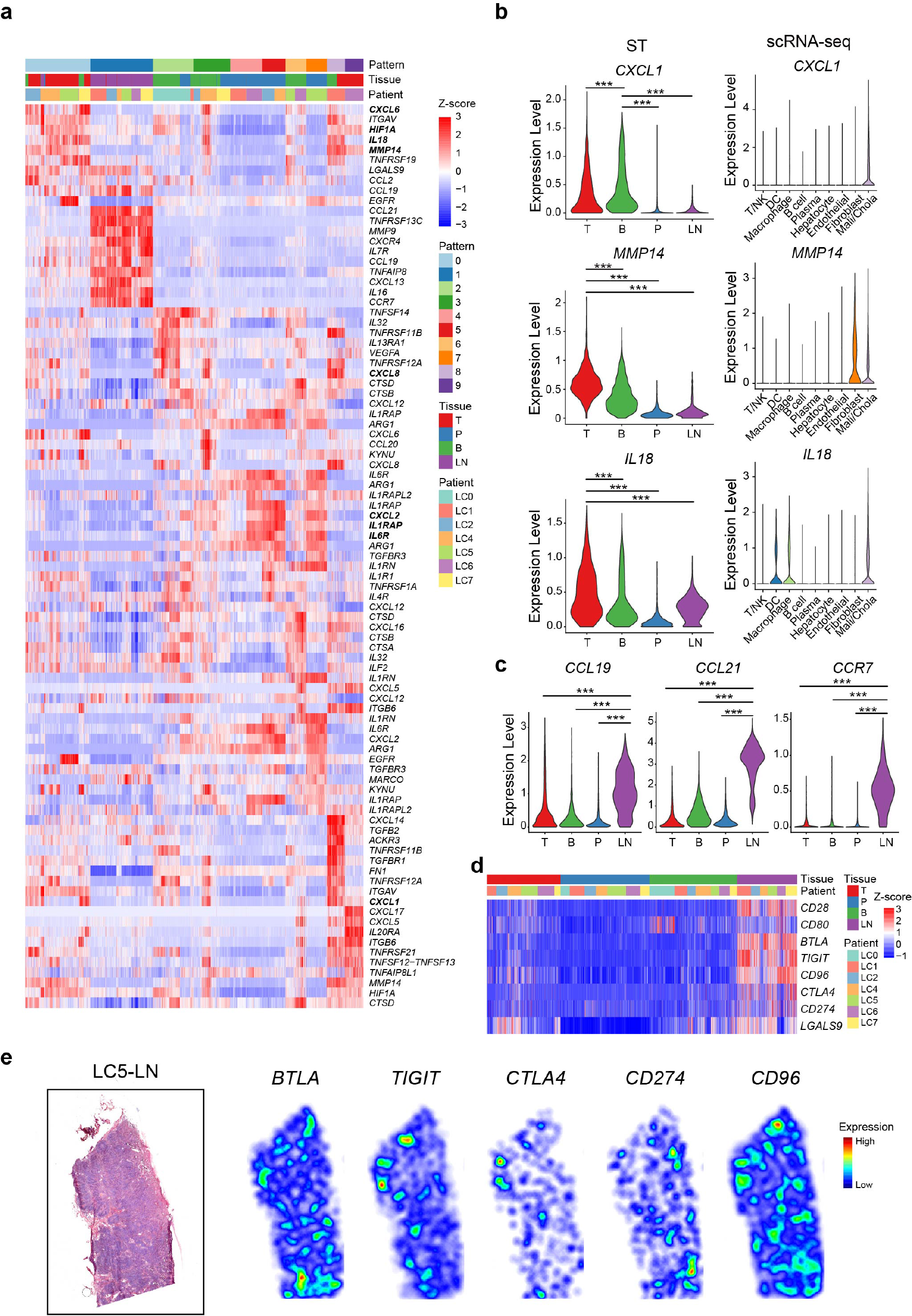
Composition and spatial heterogeneity of extracellular matrix (ECM) components in four regional sites of patients with ICC. **(a)** Heat map showing the clustering of ECM components according to differentially-expressed genes in segmented spots. Different patterns, tissues and patients are labeled by colors. **(b)** Violin plots representing the expressions of *CXCL1*, *MMP14*, and *IL18* in four regional sites by ST data (left panel) and in different cell types by scRNA-seq (right panel). **(c)** Violin plots showing the expression levels of *CCL19*, *CCL21*, and *CCR7* in four regional sites. **(d)** Heat map showing the expression levels of *CD28, CD80, BTLA, TIGIT, CD96, CTLA4, CD274*, and *LGALS9* in segmented ST spots. Tissues and patients are labeled by colors. **(e)** H&E staining and heat maps of the spatial expression of inhibitory related genes *BTLA*, *TIGIT*, *CTLA4*, *CD96*, and *CD274* in corresponding slide of LC5-LN.

**Extended Data Fig. 4.**
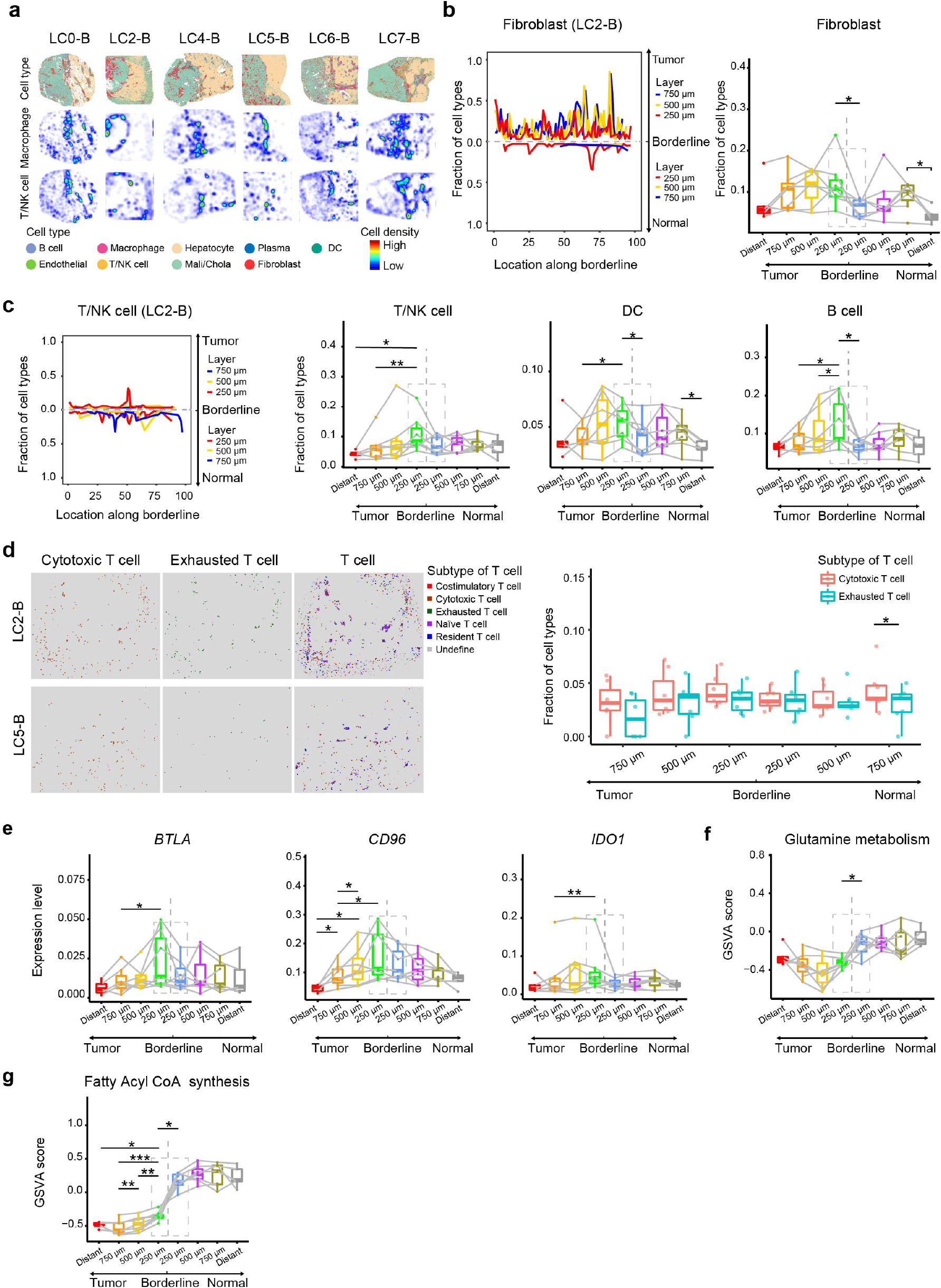
Cell states and the immune microenvironment at invasive fronts. **(a)** Spatial distribution maps of all cell types and spatial heat maps of macrophages and T/NK cells in margin areas (LC0-B, LC2-B, LC4-B, LC5-B, LC6-B, and LC7-B). **(b)** Left panel: line graphs showing the fractions of fibroblasts in all cell components along the borderline (100 subregions for each layer) of three layers from the normal and tumor sides (LC2-B). Right panel: box plots showing the fractions of fibroblasts in all cell components in different layers of margin areas. “Distant” was defined as a 250 µm wide zone in tumor or adjacent normal tissues at least 2 mm from the borderline. **(c)** Line graphs showing the fractions of T/NK cells in all cell components along the borderline (100 subregions for each layer) in three layers from the normal tumor sides (LC2-B). Box plots showing the fractions of T/NK cells, DCs, and B cells in all cell components in different layers of margin areas. **(d)** Left panel: spatial distribution maps of T cell subtypes, including co-stimulatory, cytotoxic, exhausted, naïve, and resident T cells. Right panel: box plots showing the fractions of cytotoxic T cells and exhausted T cells in all cell components in different layers of margin areas (*n* = 6). **(e)** Box plots showing the expression levels of the immune checkpoint genes *BTLA*, *CD96* and *IDO1* in different layers of margin areas (*n* = 7). **(f-g)** Box plots showing the GSVA scores of glutamine metabolism (**f**) and fatty acyl CoA synthesis (**g**) of tumor cells at the tumor side and all cell components in the paratumor side of the borderline (*n* = 7). Two-tailed paired *t*-tests were used for panel **b-g**.

**Extended Data Fig. 5.**
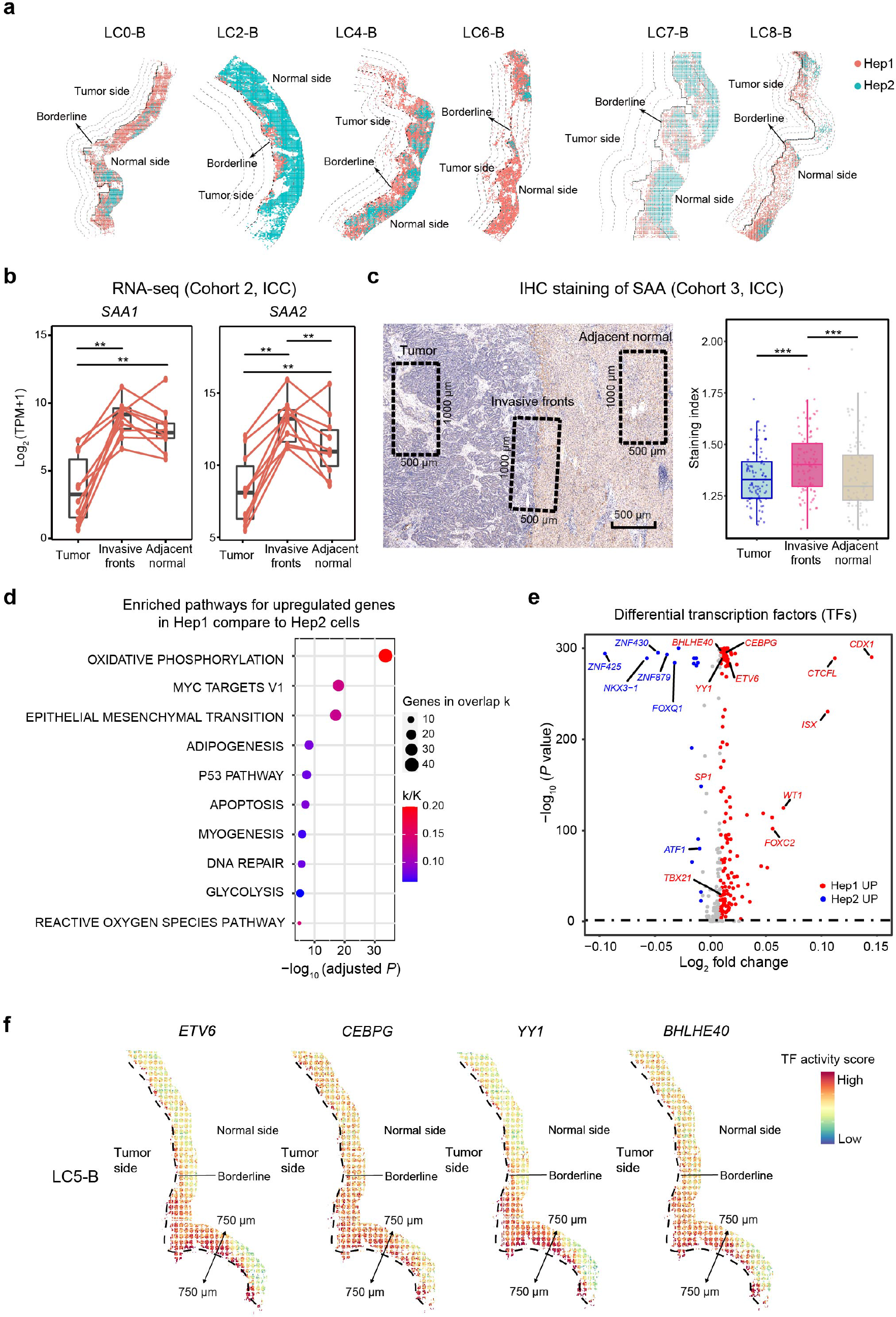
Upregulated expression of SAA in hepatocytes in invasive fronts and potential transcriptional factors (TFs). **(a)** Spatial distribution maps of Hep1 and Hep2 cells in three layers at the paratumor sides of invasive fronts of ST sides (LC0-B, LC2-B, LC4-B, LC6-B, LC7-B, and LC8-B). **(b)** *SAA1* and *SAA2* expressions of tumor tissues, the border area (a 1 cm-wide zone centered on the borderline) and adjacent normal tissues of 10 ICC patients from Validation Cohort 2 using bulk-RNA sequencing. **(c)** Diagram of the sampling zones (500 µm × 1,000 µm zone) of tumor tissues, invasive fronts, and adjacent normal tissues in immunohistochemistry (IHC) staining of SAA in ICC and analysis of the IHC staining index of SAA in the three zones of 93 ICC patients from Validation Cohort 3. Zones from tumor or adjacent normal tissues were acquired from the areas at least 1 mm from the borderline, and three different repeated zones were used to acquire an average value of the staining index. **(d)** Hallmark pathway analysis for upregulated genes in Hep1 cells compared to Hep2 cells in invasive fronts of ST sides. **(d)** Volcano plot of differentially-expressed TFs between Hep1 and Hep2 cells. The red dots represent TFs upregulated in Hep1 cells, and the blue dots represent TFs upregulated in Hep2 cells (LC5-B). **(f)** The TF activity scores of ETV6, CEBPG, YY1, and BHLHE40 in hepatocytes at invasive fronts in ST sides (LC5-B).

**Extended Data Fig. 6.**
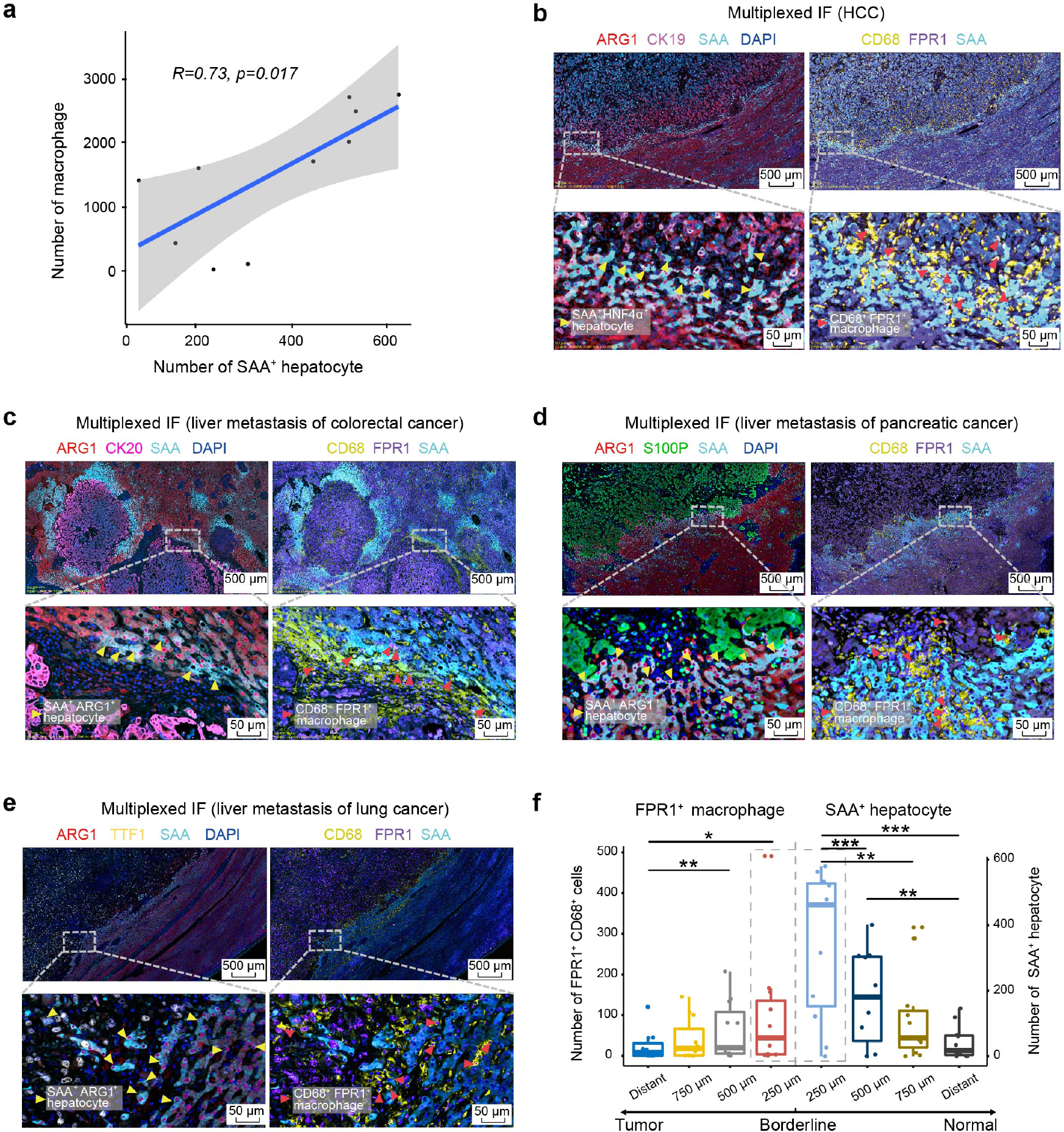
Accumulation of macrophages close to SAA^high^ hepatocytes at invasive fronts of liver cancers. **(a)** Scatter plot illustrating the correlations between the number macrophages and the number of SAA^+^ hepatocytes at invasive fronts (1,000 µm in axial length) of 10 ICC patients from Validation Cohort 1 using multiplexed IF staining. **(b)** Multiplexed IF staining (ARG1, CK19, FRP1, CD68, SAA, and DAPI) showing co-aggregation of FPR1^high^ macrophages (FPR1^+^CD68^+^ cells) and SAA^high^ hepatocytes (SAA^+^ARG1^+^ cells) at invasive fronts of HCC patients from Validation Cohort 1. **(c)** Multiplexed IF staining (ARG1, CK20, FRP1, CD68, SAA, and DAPI) showing co-aggregation of FPR1^high^ macrophages (FPR1^+^CD68^+^ cells) and SAA^high^ hepatocytes (SAA^+^ARG1^+^ cells) at invasive fronts of patients with liver metastasis of colorectal cancers from Validation Cohort 1. **(d)** Multiplexed IF staining (ARG1, S100P, FRP1, CD68, SAA, and DAPI) of FPR1^high^ macrophages (FPR1^+^CD68^+^ cells) and SAA^high^ hepatocytes (SAA^+^ARG1^+^ cells) at invasive fronts of patients with liver metastasis of pancreatic cancers from Validation Cohort 1. **(e)** Multiplexed IF staining (ARG1, TTF1, FRP1, CD68, SAA, and DAPI) of FPR1^high^ macrophages (FPR1^+^CD68^+^ cells) and SAA^high^ hepatocytes (SAA^+^ARG1^+^ cells) at invasive fronts of the patient with liver metastasis of lung cancer from Validation Cohort 1. **(f)** Cell number of macrophages (CD68^+^ cells) and the SAA^+^ hepatocytes (SAA^+^ARG1^+^ cells) in different layers (1,000 µm in axial length) at margin areas and distant sites of 11 patients with liver cancers. including HCC (*n* = 5), liver metastasis of colorectal cancer (*n* = 3), liver metastasis of pancreatic cancer (*n* = 2), and liver metastasis of lung cancer (*n* = 1) in Validation Cohort 1 using multiplexed IF staining.

**Extended Data Fig. 7.**
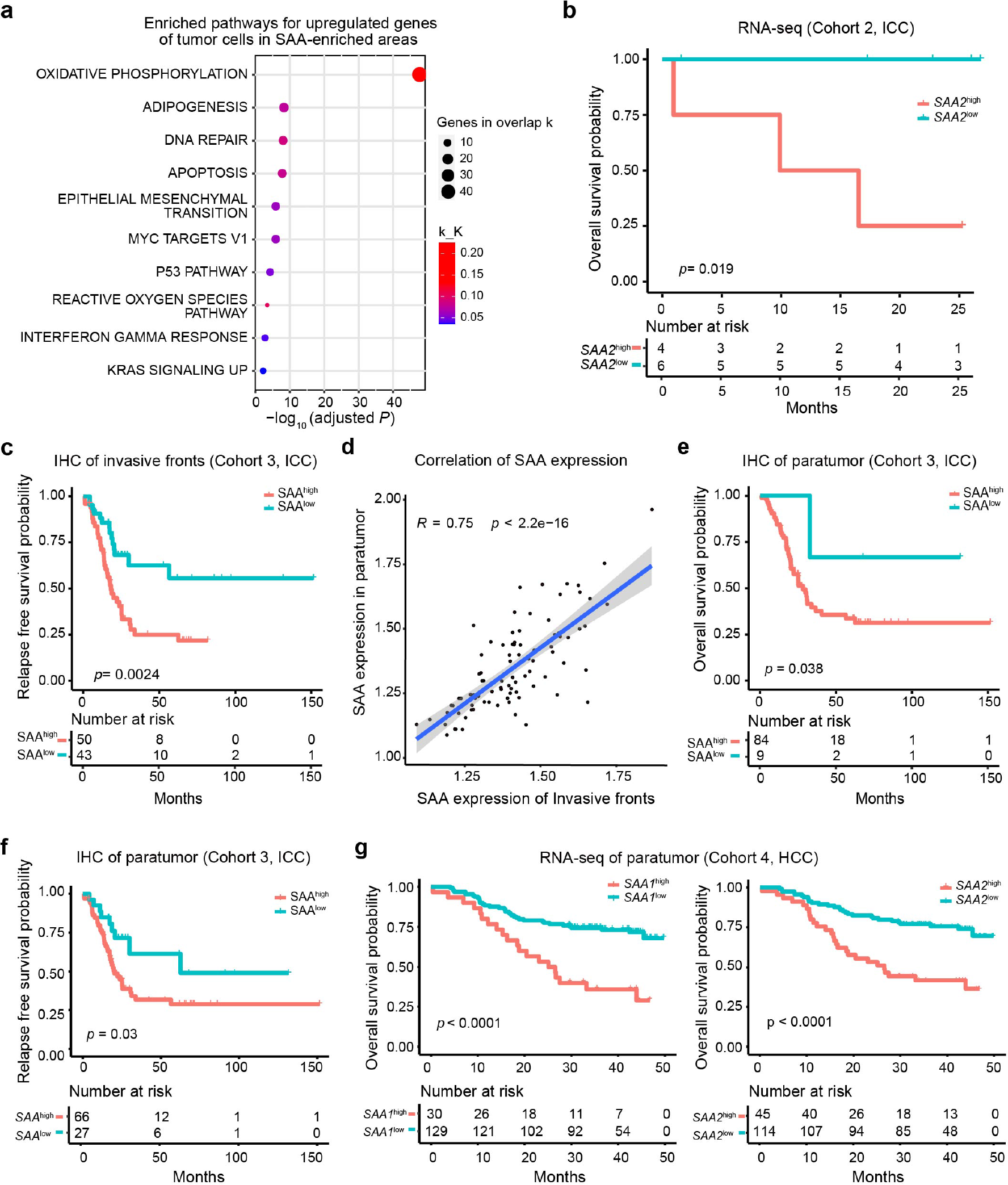
Overexpression of Serum Amyloid A (SAA) in hepatocytes at invasive fronts or paratumor tissues correlated with poor prognoses of patients with liver cancers. **(a)** Enriched pathways in hallmark genesets for upregulated genes of tumor cells in SAA-enriched areas at invasive fronts, when compared with those in remanent regions at invasive fronts in ST sides. **(b)** Overall survival (OS) curves of 10 patients with ICC from Validation Cohort 2 grouped by *SAA2* expressions in the border areas by bulk-RNA sequencing. **(c)** Relapse-free survival (RFS) curves of 93 patients with ICC from Validation Cohort 3, grouped by relative expressions of SAA at invasive fronts by IHC staining. **(d)** Scatter diagram showing the correlations between SAA expressions at invasive fronts and adjacent normal tissues by IHC staining of 93 patients with ICC from Validation Cohort 3. **(e)** OS curves of 93 patients with ICC from Validation Cohort 3, grouped by relative expressions of SAA in adjacent normal tissues by IHC staining. **(f)** RFS curves of 93 patients with ICC from Validation Cohort 3, grouped by relative expressions of SAA in adjacent normal tissues by IHC staining. **(g)** OS curves of 159 patients with HCC from Validation Cohort 4, grouped by relative expressions of *SAA1* and *SAA2* in paratumor tissues by RNA-seq data, respectively.

**Supplementary Table 1.** The clinical and pathological information of 8 ICC patients in Discovery Cohort.

| Patient ID | Sex | Age | Tumor size (cm) | lymph node metastasis | distant metastasis | AJCC stage |
| --- | --- | --- | --- | --- | --- | --- |
| LC0 | Female | 66 | $8.5 \times 8 \times 7.5$ | 0 | 0 | II |
| LC1 | Female | 65 | $4 \times 4 \times 2.5$ | 0 | 1 | IV |
| LC2 | Female | 50 | $5 \times 5 \times 5$ | 0 | 0 | Ia |
| LC4 | Male | 37 | $6.5 \times 5 \times 4$ | 1 | 0 | IIIB |
| LC5 | Female | 67 | $7 \times 3 \times 3$ | 0 | 0 | II |
| LC6 | Male | 63 | $7 \times 7 \times 5$ | 1 | 0 | IIIB |
| LC7 | Male | 65 | $2.1 \times 1.8 \times 1.5$ | 0 | 0 | Ia |
| LC8 | Female | 82 | $3.5 \times 3 \times 2$ | 0 | 0 | II |

**Supplementary Table 2.** Detailed information of spatial transcriptomics slides of samples from 8 ICC patients.

| **Patients ID** | **Tumor (T)** | **Adjacent**<br>**normal (P)** | **Margin area**<br>**(B)** | **Lymph Node (LN)** |
| --- | --- | --- | --- | --- |
| LC0 | 0 | 2 | 3 | 0 |
| LC1 | 2 | 2 | 2 | 2 |
| LC2 | 2 | 2 | 1 | 2 |
| LC4 | 2 | 2 | 2 | 1 |
| LC5 | 2 | 2 | 1 | 2 |
| LC6 | 2 | 2 | 1 | 2 |
| LC7 | 2 | 2 | 1 | 2 |
| LC8 | 0 | 0 | 2 | 0 |

**Supplementary Table 3.**
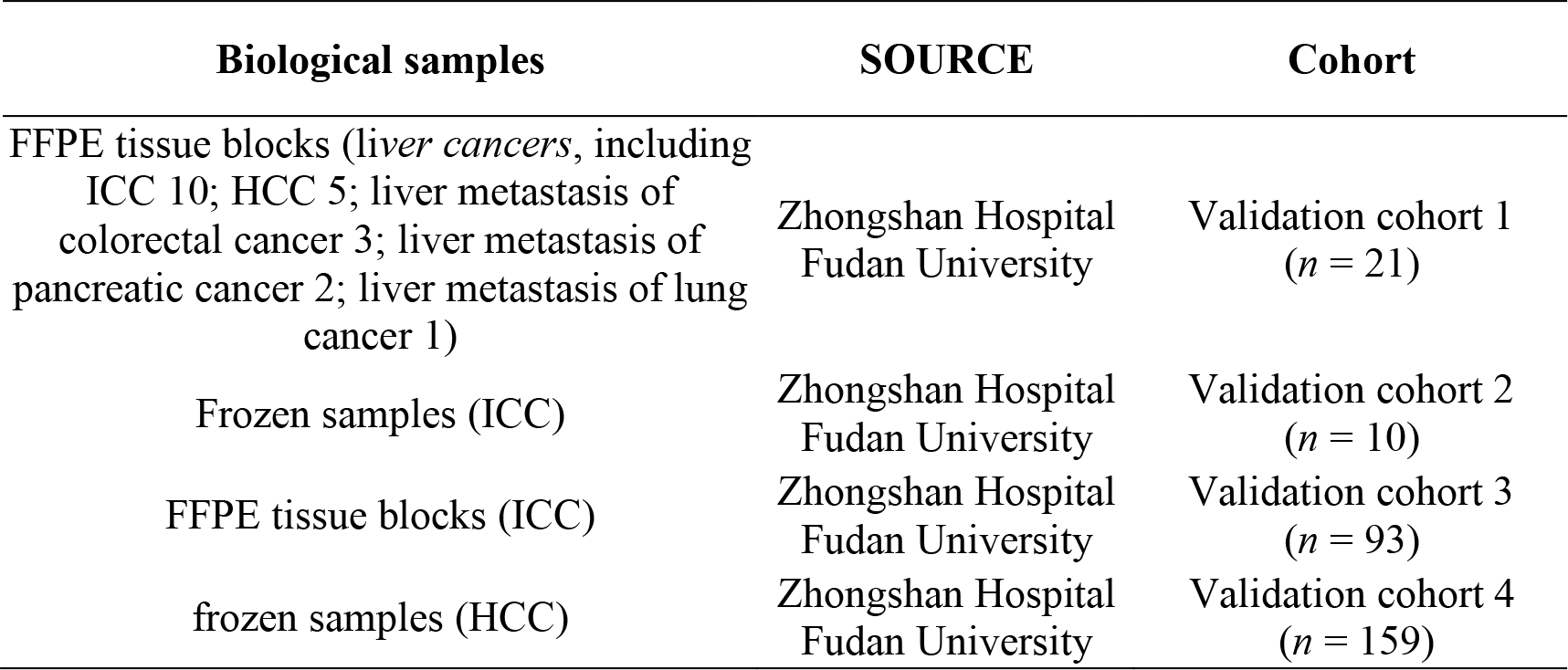
Study subjects resource and cohorts.

| Biological samples | SOURCE | Cohort |
| --- | --- | --- |
| FFPE tissue blocks (*liver cancers*, including ICC 10; HCC 5; liver metastasis of colorectal cancer 3; liver metastasis of pancreatic cancer 2; liver metastasis of lung cancer 1) | Zhongshan Hospital<br>Fudan University | Validation cohort 1<br>(*n* = 21) |
| Frozen samples (ICC) | Zhongshan Hospital<br>Fudan University | Validation cohort 2<br>(*n* = 10) |
| FFPE tissue blocks (ICC) | Zhongshan Hospital<br>Fudan University | Validation cohort 3<br>(*n* = 93) |
| frozen samples (HCC) | Zhongshan Hospital<br>Fudan University | Validation cohort 4<br>(*n* = 159) |

**Supplementary Table 4.**
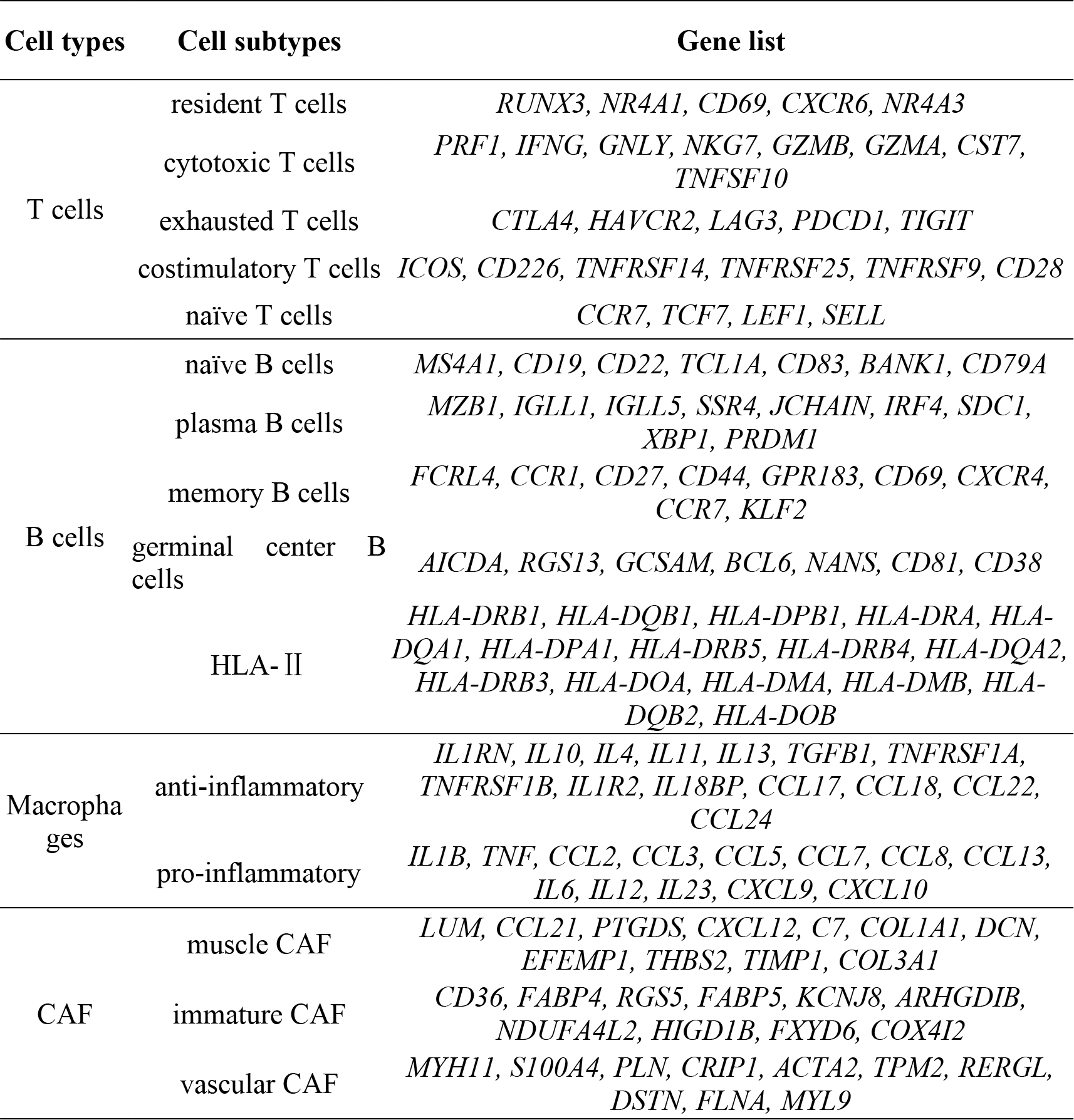
Selected marker genes in different cell subtypes.

## Notes

### Competing Interest Statement

The authors have declared no competing interest.

